# mRNA-based generation of marmoset PGCLCs capable of differentiation into gonocyte-like cells

**DOI:** 10.1101/2022.09.20.508677

**Authors:** Musashi Kubiura-Ichimaru, Christopher Penfold, Kazuaki Kojima, Constance Dollet, Haruka Yabukami, Katsunori Semi, Yasuhiro Takashima, Thorsten Boroviak, Hideya Kawaji, Knut Woltjen, Aki Minoda, Erika Sasaki, Toshiaki Watanabe

## Abstract

Primate germ cell development remains largely unexplored due to limitations in sample collection and the long duration of development. In mice, primordial germ cell-like cells (PGCLCs) derived from pluripotent stem cells (PSCs) can develop into functional gametes by *in vitro* culture or *in vivo* transplantation. Such PGCLC-mediated induction of mature gametes in primates is highly useful for understanding human germ cell development. Since marmosets generate functional sperm earlier than other species, recapitulating the whole male germ cell development process is technically more feasible. Here, we induced the differentiation of iPSCs into gonocyte-like cells via PGCLCs in marmosets. First, we developed an mRNA transfection-based method to efficiently generate PGCLC. Subsequently, to promote PGCLC differentiation, xenoreconstituted testes (xrtestes) were generated in the mouse kidney capsule. PGCLCs show progressive DNA demethylation and stepwise expression of developmental marker genes. This study provides an efficient platform for the study of marmoset germ cell development.

**Highlights:**

1. An mRNA transfection-based PGCLC induction method is developed
2. Marmoset PGCLCs are efficiently induced from iPSCs
3. Marmoset PGCLCs differentiate into gonocyte-like cells in mouse kidneys
4. Developmentally regulated expression and demethylation are recapitulated

**eTOC blurb:** Watanabe et al. efficiently induced marmoset primordial germ cell-like cells (PGCLCs) using an mRNA transfection-based approach. PGCLCs further develop into gonocyte-like cells in the xenoreconstituted testes constructed under mouse kidney capsules. This system faithfully reproduced *in vivo* developmental processes (e.g., stage-specific expression of developmentally regulated genes and DNA demethylation).

## Introduction

The testis is known to be among the fastest evolving tissue in terms of change to gene expression (Brawand et al., 2011; Khaitovich et al., 2006; Swanson and Vacquier, 2002). This is due to the selection pressure acting on spermatogenic cells, only a small portion of which leads to fertilization. Despite large differences across species, both in terms of embryo development and germ cell-specific transcriptional programs, studies of mammalian spermatogenic cells have been largely restricted to mice. Therefore, developing appropriate primate models and understanding the mechanisms of primate germ cell development is paramount of importance for the treatment of infertility arising from gametogenesis defects.

Marmosets (*Callithrix jacchus*) are new world monkeys native to Brazil that are often used in biomedical research, particularly in brain science, owing to their small size, relative ease of handling, high reproductive ability, and close evolutionary relationship to humans. In addition to these characteristics, marmosets reach puberty within 1 year of birth. This relatively short period of sexual maturation makes marmosets an ideal model organism for studying primate germ cell development. However, the high cost of studies using marmosets limits the number of animal experiments. Therefore, to complement *in vivo* studies, it is important to develop a tractable marmoset germ cell developmental system using pluripotent stem cells (PSCs).

Several studies have reported the induction of primordial germ cell-like cells (PGCLCs) from PSCs in primates (humans, macaques, and marmosets)(Irie et al., 2015; Sakai et al., 2020; Sasaki et al., 2015; Sosa et al., 2018; Yoshimatsu et al., 2021). Although complete differentiation of PGCLCs into sperm or eggs has been successful in mice, it has not yet been achieved in any primate species. In humans, PGCLCs differentiate into gonocytes after a few months of the *in vitro* culture of xenoreconstituted testes (xrtestes) generated from human PGCLCs and mouse embryonic gonadal somatic cells (Hwang et al., 2020). Furthermore, in rhesus macaques, male PGCLCs differentiate into MAGEA4-positive gonocytes by homologous transplantation into adult testes and xenotransplantation into seminiferous tubules of mouse testes (Sosa *et al*., 2018). Because both extrinsic and intrinsic factors determine developmental speed, the duration of germ cell development in the PGCLC system is likely to be influenced by the *in vivo* development timetable. Although early embryonic development is delayed in marmosets (Phillips, 1976), they reach sexual maturation earlier than other primates. Therefore, the marmoset PGCLC system may be more feasible for recapitulating the entire primate germ cell development process. Although PGCLCs have been generated in marmosets, the reported method requires *a priori* transgene integration for the forced expression of *SOX17* and *PRDM1* (Yoshimatsu et al., 2021). Furthermore, there are no reports on the differentiation of PGCLCs into MAGEA4-positive gonocytes in marmosets.

In addition to examining germ cell development, this PGCLC-initiated germ cell developmental system is useful for generating genetically modified animals. This is enabled by gamete production using this system from genetically modified PSCs. All genetically modified marmosets generated to date have been produced from zygotes (Park and Sasaki, 2020). Cultured cell-based systems, like PGCLC-mediated system, enable the creation of animals with complex genetic modifications such as reporter gene knock-in and multiple modifications. Furthermore, a cell-based system ensures the production of the expected genetic modifications. Thus, establishing a germ cell developmental system from PGCLCs to produce functional gametes may accelerate primate research.

In this study, to develop a PGCLC-initiated germ cell developmental system in marmosets, we developed a novel mRNA transfection-based method to convert PSCs into PGCLCs. Furthermore, these PGCLCs were differentiated into a gonocyte-like state using a transplantation approach. Our results provide the basis for studying marmoset gametogenesis using PGCLCs.

## Results

### mRNA transfection-based induction of marmoset PGCLCs

We have previously reported mRNA transfection-based methods for marmoset iPS cell induction (Watanabe et al., 2019). To generate PGCLCs from iPSCs, we performed mRNA transfection. Because *SOX17* has been reported to have critical functions in PGCLC induction in humans (Irie *et al*., 2015; Kobayashi et al., 2017), we selected this mRNA for transfection. To alleviate the damage caused by mRNA transfection in cells, we transfected cells with interferon suppressors (vaccinia virus *E3*, *K3*, and *B18R* mRNAs)(Poleganov et al., 2015) and an apoptosis suppressor (mouse P53DD mRNA)(Hong et al., 2009). To monitor the differentiation state, IRES-mCherry was knocked into the 3′-UTR of marmoset *NANOS3*, a highly conserved germline-specific gene (Figure 1B). This genetic modifications were performed using an female iPSC line (mRNA iPSC) previously induced in adult liver cells (Watanabe *et al*., 2019).

**Figure 1.**
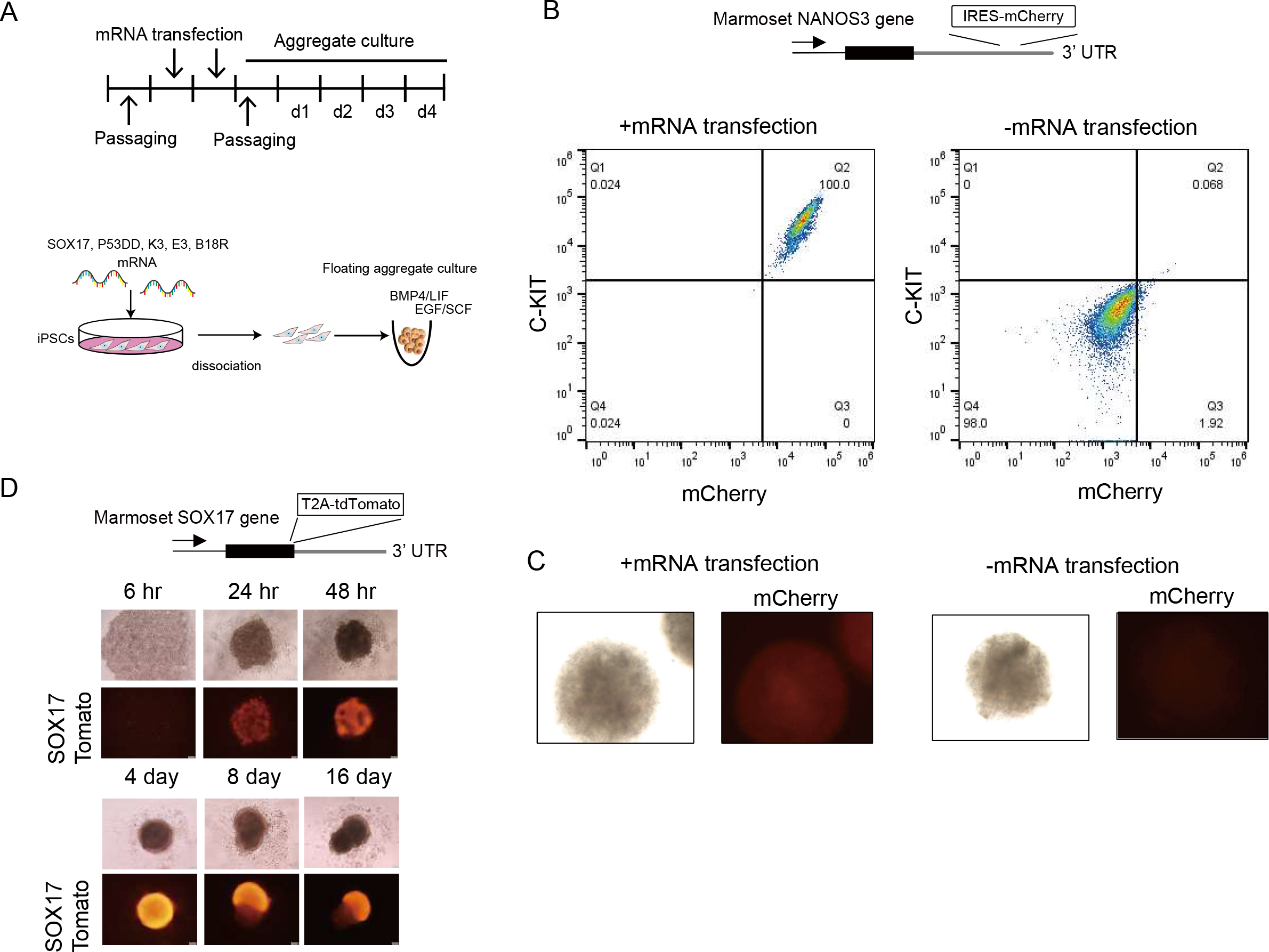
mRNA-based generation of PGCLCs from iPSCs. **A** Schematic diagram for the generation of PGCLCs from iPSCs. **B** mRNA-mediated induction of PGCLCs. An IRES-mCherry cassette was inserted into the 3′-UTR of the *NANOS3* gene to monitor differentiation into PGCLCs. The two charts represent the results of the normal induction procedure (left) or mRNA transfection step omission from it (right). **C** Microscopic observation of mCherry fluorescence of PGCLCs. PGCLCs were generated using the same conditions as B. **D** Time course examination of *SOX17*-tdTomato fluorescence.

After two successive days of transfection, the transfected cells were seeded onto low-attachment 96-well plates to form aggregates in medium containing LIF, EGF, SCF, and BMP4 based on the procedures for human PGCLC induction (Figure 1A)(Hwang *et al*., 2020; Irie *et al*., 2015; Kojima et al., 2017; Sasaki *et al*., 2015; Yamashiro et al., 2018). Four days after making the aggregate, C-KIT expression was observed in many cells (>80%) in the aggregates (Figure 1B). These C-KIT-positive cells exhibited a *NANOS3*-mCherry signal. In contrast, when mRNA transfection was omitted, neither C-KIT nor *NANOS3*-mCherry expression was observed (Figure 1B), indicating that mRNA transfection plays a critical role in PGCLC induction.

Since the *NANOS3*-mCherry signal was weak for microscopic observation (Figure 1C). We inserted T2A-tdTomato into the C-terminus of *SOX17* gene (*SOX17*-tdTomato (ST)) of two male iPS lines harboring CAG-EGFP (CE) transgenes (971 STCE and 972 STCE). Both iPSCs were originally derived from ear fibroblasts, and our mRNA transfection-based method converted these iPSCs into PGCLCs with high efficiency (>80%) (Figure 1D and S1). Since ST showed stronger fluorescent signals (Figure 1D and S1), we used this transgene for the following experiments.

To examine the time-course change, aggregate culture was continued with medium change every four days using a male iPS line harboring *SOX17*-tdTomato transgene (971 STCE). ST expression was first observed 24 h after PGCLC aggregate formation (Figure 1D). Initially, most cells expressed ST fluorescence; however, the number of cells negative for fluorescence gradually increased over time (Figure 1D). Despite this increase in negative cells, a significant portion of cells still expressed ST, even on day 16. Thus, PGCLCs can be maintained in aggregate culture for at least a few weeks.

The optimal number of mRNA transfections was determined to be two. Two successive days of transfections resulted in higher *NANOS3* expression in day 4 aggregates (d4_PGCLC) than did single transfection, but three successive day transfections did not result in increased *NANOS3* expression (Figure S2A). The optimal number of transfected cells was determined to be 50,000 cells per well in a 6-well plate (Figure S2B). Using an increased number of cells (100,000 cells) resulted in a decreased PGCLC induction efficiency, likely due to an insufficient number of mRNAs distributed into each cell when a larger quantity was used.

### Single-cell analysis of iPSCs and PGCLCs

To examine whether induced cells showed gene expression typical of PGCs, single-cell (sc) RNA-seq libraries were generated from iPSCs (mRNA iPSCs) and d4_PGCLC aggregates (derived from mRNA iPSCs). Principle component analysis (PCA) clearly separated the iPSC and PGCLC populations (Figure 2A), which were located on opposite sides of PC1. Well-characterized iPSC (*SOX2* and *ZIC2*) and PGC markers (*SOX17*, *NANOS3*, *KIT*, and *DND1*) were identified in the genes contributing to PC1 (Figure 2B). This suggests the successful generation of PGCLCs. In addition, this analysis revealed the presence of mesoendoderm-like cells (*NODAL* and *MIXL1*), a small number of endoderm-like cells (*SOX17* and *FOXA2*), mesenchymal-like cells (*HAND1*, *ANXA1*, and *VIM*), and amnion-like cells (*HAND1*, *TFA2PC*, and *GABRP*). Although mesoendoderm-like cells were found only in the iPSC library, amnion-like cells were found only in the PGCLC library (Figure 2C).

**Figure 2.**
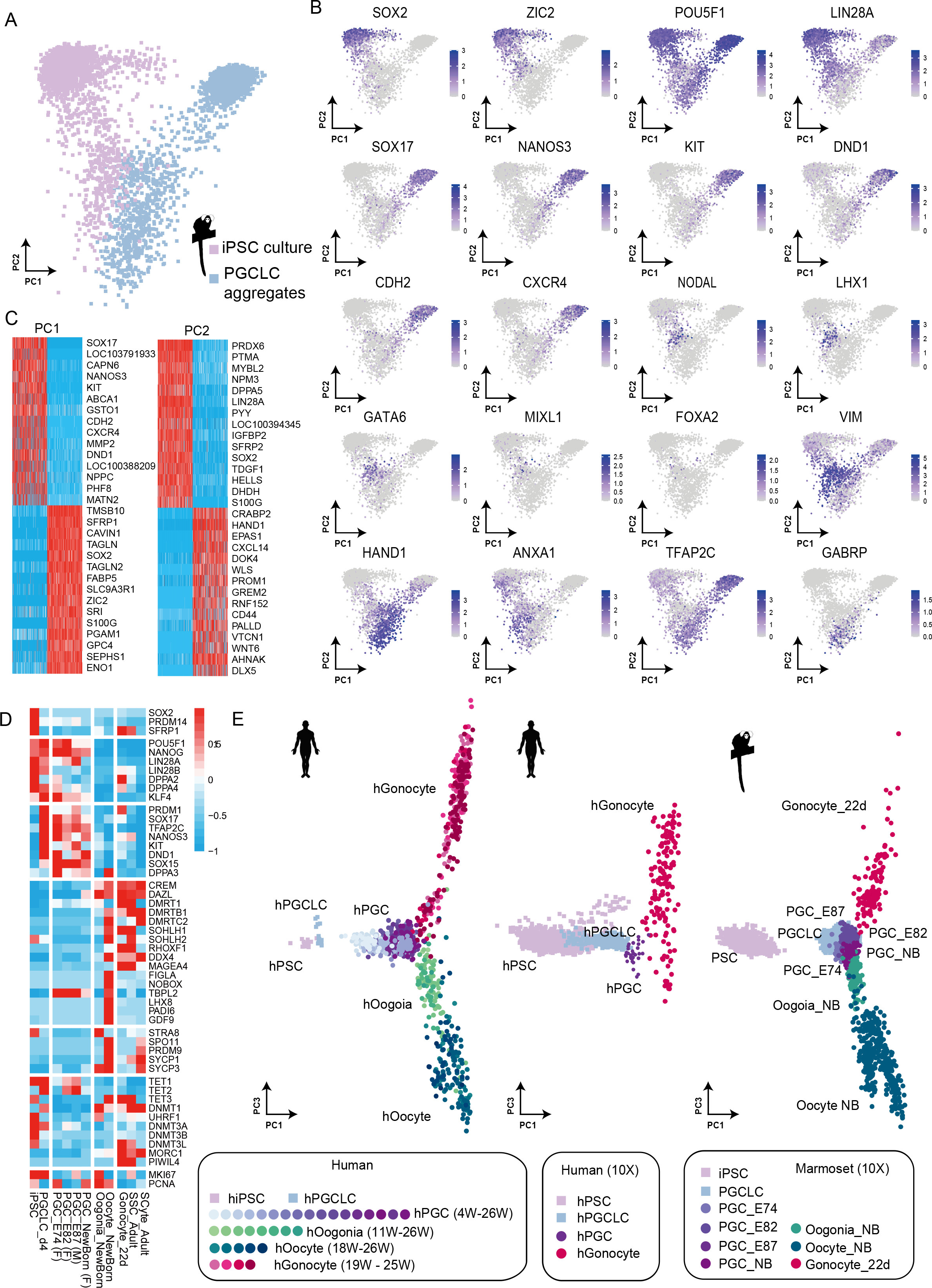
Gene expression analyses of marmoset PGCLCs induced from iPSCs. **A** Single-cell analyses of iPSCs and PGCLCs. **B** Well-characterized iPSC markers (*SOX2* and *ZIC2*) and PGC markers (*SOX17, NANOS3, KIT*, and *DND1*) were found in the genes contributing to PC1. **C** In addition to the iPSCs and PGCLCs, mesoendoderm-like cells (*NODAL* and *MIXL1*), a small number of endoderm-like cells (*SOX17* and *FOXA2*), mesenchymal-like cells (*HAND1, ANXA1*, and *VIM*), and amnion-like cells (*HAND1, TFA2PC*, and *GABRP*) were present. **D** Heatmap analyses of marker genes. iPSCs, PGCLCs, and marmoset endogenous developmental germ cells were examined. **E** PCA of PSCs, PGCLCs, and developing germ cells from humans (left and center) and marmosets (right). Published human data were downloaded(Chen *et al*., 2019; Kojima *et al*., 2017; Li *et al*., 2017; Sohni *et al*., 2019) and analyzed together with marmoset data. Human data are shown in two separate panels according to the scRNA-seq platform. The anchoring function was used to integrate datasets of different platforms.

### Single-cell analysis of developing gonads

Before setting out to advance the development of PGCLCs, we collected information on *in vivo* developing germ cells as a reference. We thus prepared scRNA-seq libraries from developing marmoset gonads. Ovaries from embryonic day 74 (E74), E82, and newborn marmosets and testes from E74, E87, 22 day-olds (22d), and 3-year and 10-month-old marmosets were subjected to single-cell analyses. After tSNE plotting, the germ cells of interest were extracted from each library (Figure S3). PGCs marked by *POU5F1* expression were extracted from fetal ovaries (E74 and E82) and testes (E87) (Figure S3). In E74 fetal testes, only a small number of PGCs were present, and they did not form a distinct cluster (Figure S3). Therefore, we did not extract the PGCs from this library. This small number is probably because many PGCs are still migrating and have not yet arrived at this stage (Aeckerle et al., 2015). In the somatic cells of this E74 testis and its sibling E74 ovary, the expression of genes involved in initial sex differentiation was observed (*SRY*, *SOX9*, *AMH* in testes, and *FOXL2* in ovaries) (Figure S4), suggesting that sex differentiation is already initiated. Consistent with a previous report (Fereydouni et al., 2014), a wide range of cells (PGCs, oogonia, and early oocytes) were found in newborn ovaries. In 22d testes, germ cells were already differentiated into gonocytes, as shown by the decreased expression of *POU5F1* and elevated expression of *DDX4*. These expression patterns in 22d testis germ cells align with previously published studies (Albert et al., 2010; McKinnell et al., 2013; Mitchell et al., 2008).

### PGCLCs align with *in vivo* PGCs

To examine whether our induced PGCLCs showed similar gene expression patterns to *in vivo* PGCs, we compared the expression of representative marker genes between *in vivo* germ cells and our PGCLCs. A similar pattern was observed between *in vivo* PGCs and PGCLCs for the key marker genes (Figure 2D). Both *in vivo* PGCs and PGCLCs showed the expression common marker genes for PSCs and PGCs (e.g., *POU5F1, NANOG, LIN28A, DPPA4*, and *KLF4*) and PGC marker genes (*PRDM1, SOX17, TFAP2C, NANOS3, KIT, DND1*, and *SOX15*). However, none of the PSC marker genes (e.g., *SOX2*), gonocyte/oogonium genes (e.g., *DAZL* and *SOHLH1*), oocyte genes (e.g., *FIGLA* and *NOBOX*), or meiotic genes (e.g., *STRA8* and *SPO11*) were expressed in both *in vivo* PGCs and PGCLCs.

To obtain further evidence that our PGCLCs were indeed PGC-like, PCA was conducted using marmoset iPSCs, PGCLCs, and *in vivo* germ cells (PGCs, gonocytes, oogonia, and oocytes). As expected, marmoset PGCLCs clustered together with *in vivo* PGCs (Figure 2E, right panel). Furthermore, we added human data to this analysis (Chen et al., 2019; Kojima *et al*., 2017; Li et al., 2017; Sohni et al., 2019) to obtain additional supporting evidence. Human PSC, PGCLCs, and germ cells from various developmental stages aligned well with their corresponding cells in marmosets (Figure 2E). These results suggest that human and marmoset early germ cell development is overall conserved and that our PGCLCs were indeed reminiscent of *in vivo* PGCs.

### Generation of xrtestes in mouse kidneys

To advance the development of marmoset PGCLCs, we used the xrtestis system that has been used to differentiate human PGCLCs (Hwang *et al*., 2020). However, instead of the *in vitro* air-liquid interface culture used in humans (Hwang *et al*., 2020; Yamashiro *et al*., 2018), we employed mouse kidney transplantation, as this system develops mouse reconstituted testes well (Matoba and Ogura, 2011). To recapitulate the male *in vivo* developmental process, we used two male iPS cell lines harboring the ST and CAG-EGFP (CE) transgenes (971-STCE and 972-STCE) (Figure S1). To prepare xrtestes, FACS-purified d4_PGCLCs were mixed with E13.5 testis somatic cells for floating aggregate culture. The next day, the aggregates were transplanted into kidney capsules (Figure 3A).

**Figure 3.**
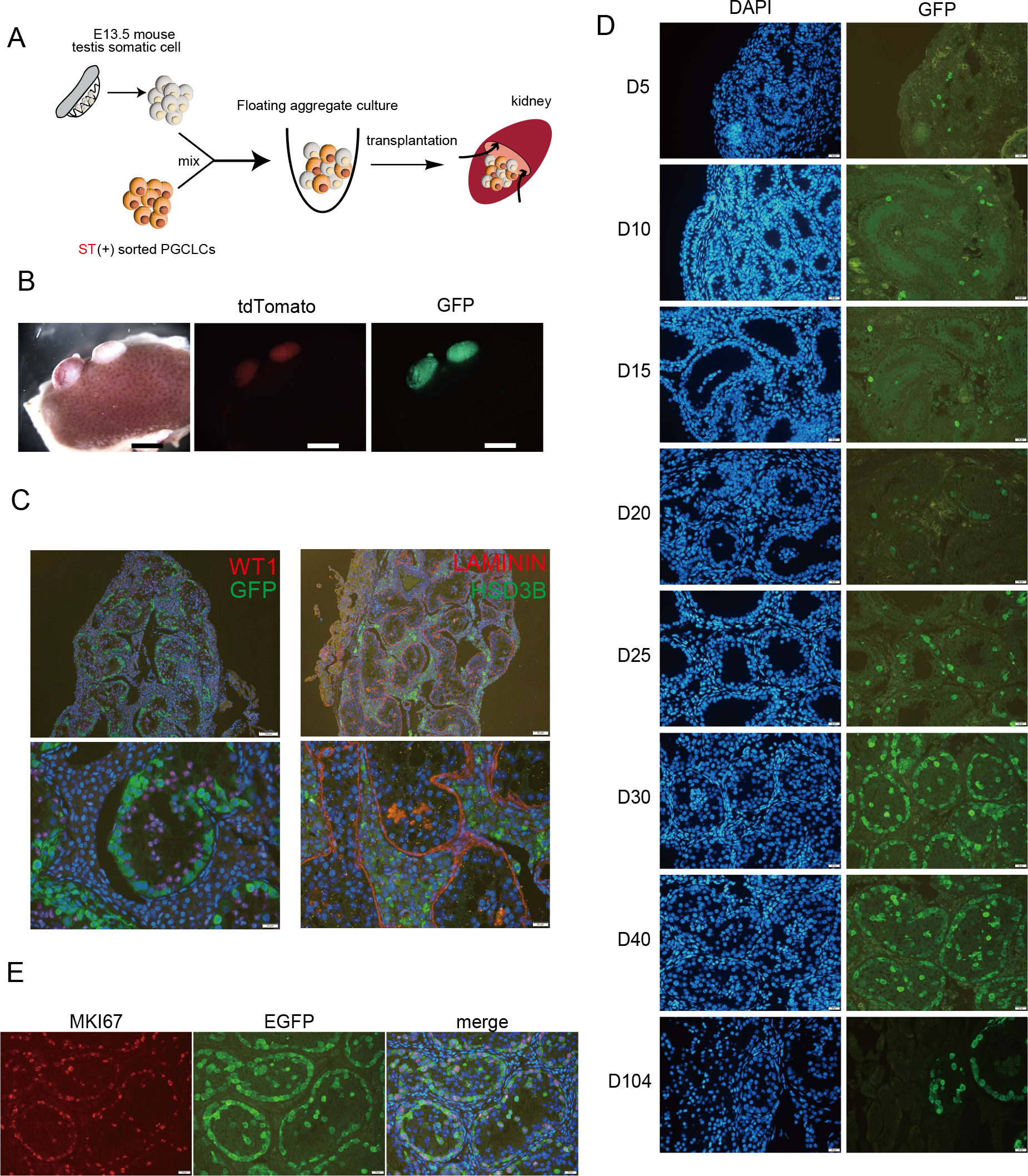
Proliferation of PGCLCs in xrtestis formed in mouse kidneys. **A** Scheme for differentiation of PGCLCs. **B** d28_xrtestes formed under the kidney capsule. PGCLC (972-STCE)-derived cells express GFP and tdTomato. Scale bar: 2 mm. **C** Reconstitution of testis structure (d104_xrtestes). Immunohistochemical analyses of the transplanted tissues. Markers: LAMININ (basement membrane), HSD3B (Leydig cells), and WT1 (Sertoli cells). PGCLC (971-STCE)-derived cells express GFP. Scale bars: 100 μm (top) and 20 μm (bottom). **D** Time course change in the number of PGCLC (971-STCE)-derived cells during xrtestis development. Days after transplantation are indicated on the left side of the pictures. Scale bar: 20 μm. **E** Many PGCLC (971-STCE)-derived cells express MKI67 in d30_xrtestes. Scale bar: 20 μm.

Ten days after transplantation (day 10), the transplanted aggregates formed a testicular cord-like structure (Figure 3B, C, D). This structure was maintained for over 100 days (Figure 3C, D) unless cancer developed. In the xrtestes, PGCLC-derived cells (EGFP- and tdTomato-positive) and Sertoli cells (WT1-positive) were found within the cord structure (marked by LAMININ). In contrast, cells expressing the Leydig cell marker HSDB were found in interstitial regions (Figure 3C). Only a small number of PGCLC-derived cells was observed in the xrtestes until day 30 after transplantation. However, by day 30, the number of PGCLC-derived cells dramatically increased, occupying the entire circumference of each tubule (Figure 3E). Consistent with this massive increase in cell number, MKI67 signals were observed in many PGCLC-derived cells from day 30 xrtestes (d30_xrtestes) (Figure 3E). Thus, PGCLCs are incorporated into the tubules of the xrtestes, and these PGCLC-derived cells actively proliferate within the tubules.

### Differentiation of PGCLCs in xrtestes

Human PGCLCs develop into gonocyte-like cells over ~80 days (Hwang *et al*., 2020). We examined the expression of four well-characterized developmentally regulated marker genes (TFAP2C, DDX4, MAGEA4, and PIWIL4) in the xrtestes from several developmental points. The results for the two iPSC lines, 972-STCE and 971-STCE, are shown in Figures 4 and S5, respectively. TFAP2C (PGC marker) expression was observed in all PGCLC-derived cells before day 40 (Figure 4A, S5). Subsequently, the number of PGCLC-derived cells expressing TFAP2C decreased dramatically. On day 81, none of the cells expressed this gene. No cells expressed gonocyte markers (DDX4, MAGEA4, and PIWIL4) on day 14. Almost all PGCLC-derived cells showed DDX4 expression on day 28 (Figure 4B and S5). On day 28, the expression of MAGEA4 was also detected in only a small number of cells. As development progressed, the proportion of cells expressing MAGEA4 increased, and all PGCLC-derived cells exhibited expression on day 81 (Figure 4C, S5). PIWIL4 was first observed in a small number of cells on day 81 (Figure 4D and S5). Thus, these analyses revealed stepwise (in)activation of gonocyte (PGC) markers during xrtestis development, and the *in vivo* developmental pattern was recapitulated in our xrtestis system(Albert *et al*., 2010; Mitchell *et al*., 2008). RT-qPCR analyses also revealed the upregulation of these and some other gonocyte-expressed genes (*CREM, DMRT1, DMRT1B, DAZL, ZBTB16, FOXR1, RHOXF1, SOHLH1, SOHLH2*, and *RBM46*) in d56, d81, and d104_xrtestes (Figure S6).

**Figure 4.**
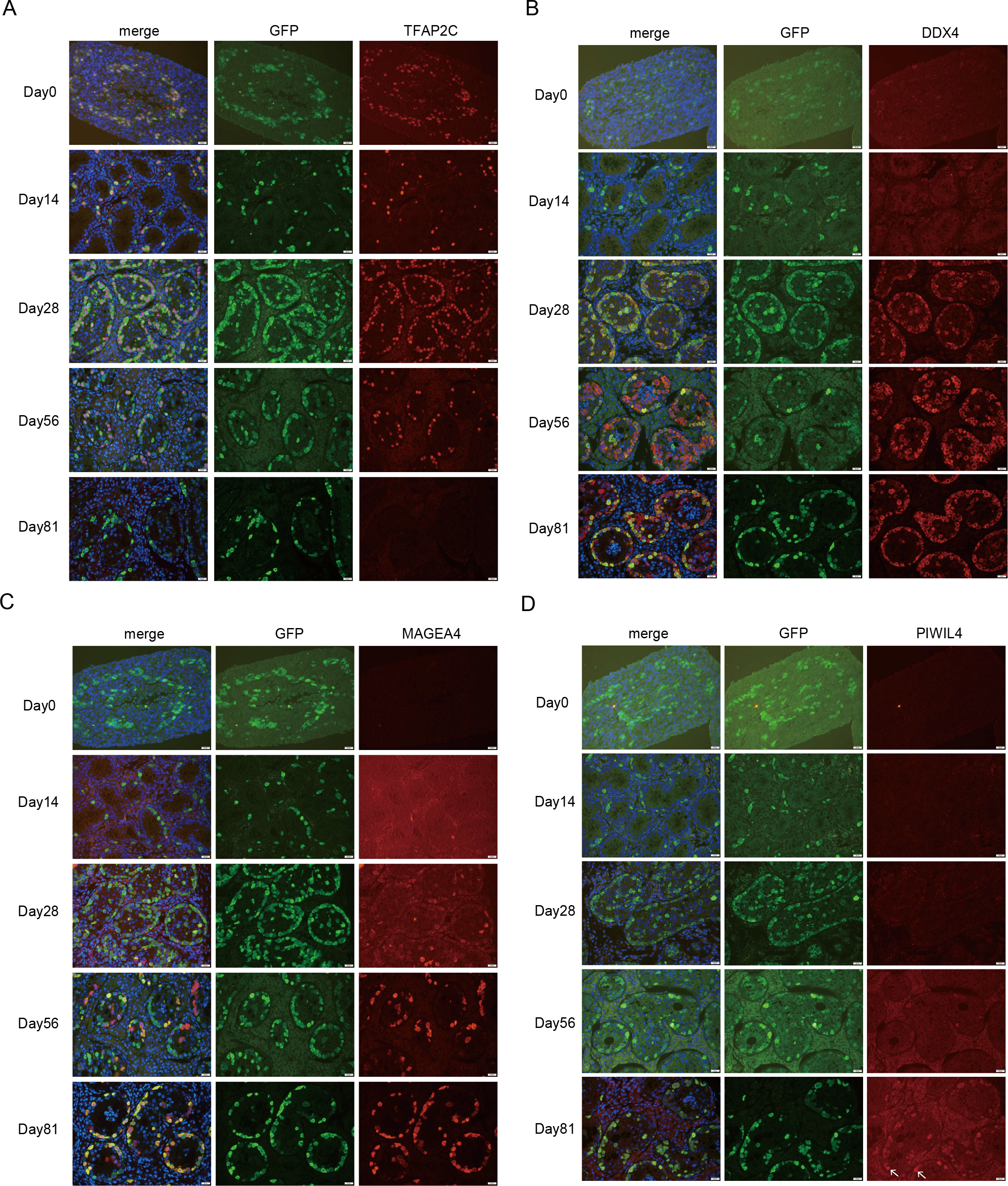
Differentiation of marmoset PGCLCs into Gonocyte-like cells. Immunofluorescence analyses of marker gene expression in developing xrtestes (972-STCE) and d5_PGCLC aggregates. TFAP2C (**A**), DDX4 (**B**), MAGEA4 (**C**), and PIWIL4 (**D**). White arrows indicate nuclear PIWIL4 staining. These staining patterns were different from those of cytoplasmic EGFP staining. Scale bar: 20 μm.

### Demethylation of PGCLCs in xenoreconstituted testes

PGC development is accompanied by progressive loss of DNA methylation (Shirane et al., 2016). To determine DNA methylation status, we conducted single-cell bisulfite-sequencing (scBS-seq) analyses in PGCLCs and PGCLC-derived cells in the xrtestes. Simultaneously, RNA expression was analyzed in the same single cells using the cytoplasmic fraction. In d4_PGCLCs, The average DNA methylation level was 61.1% (Figure 5A). Interestingly, the level decreased to 45.7% in d12_PGCLCs, suggesting the occurrence of demethylation during long floating aggregate culture. The xrtestes were generated using d4_PGCLCs. DNA methylation levels decreased gradually in xrtestes (Figure 5A). Although we still detected some residual methylation in d30_xrtestes (9.4%), this level is close to the minimum level in d104_xrtestes (4.3%). Thus, DNA demethylation was recapitulated in our xrtestis system. However, the establishment of DNA methylation was likely not initiated, even in d104_xrtestes.

**Figure 5.**
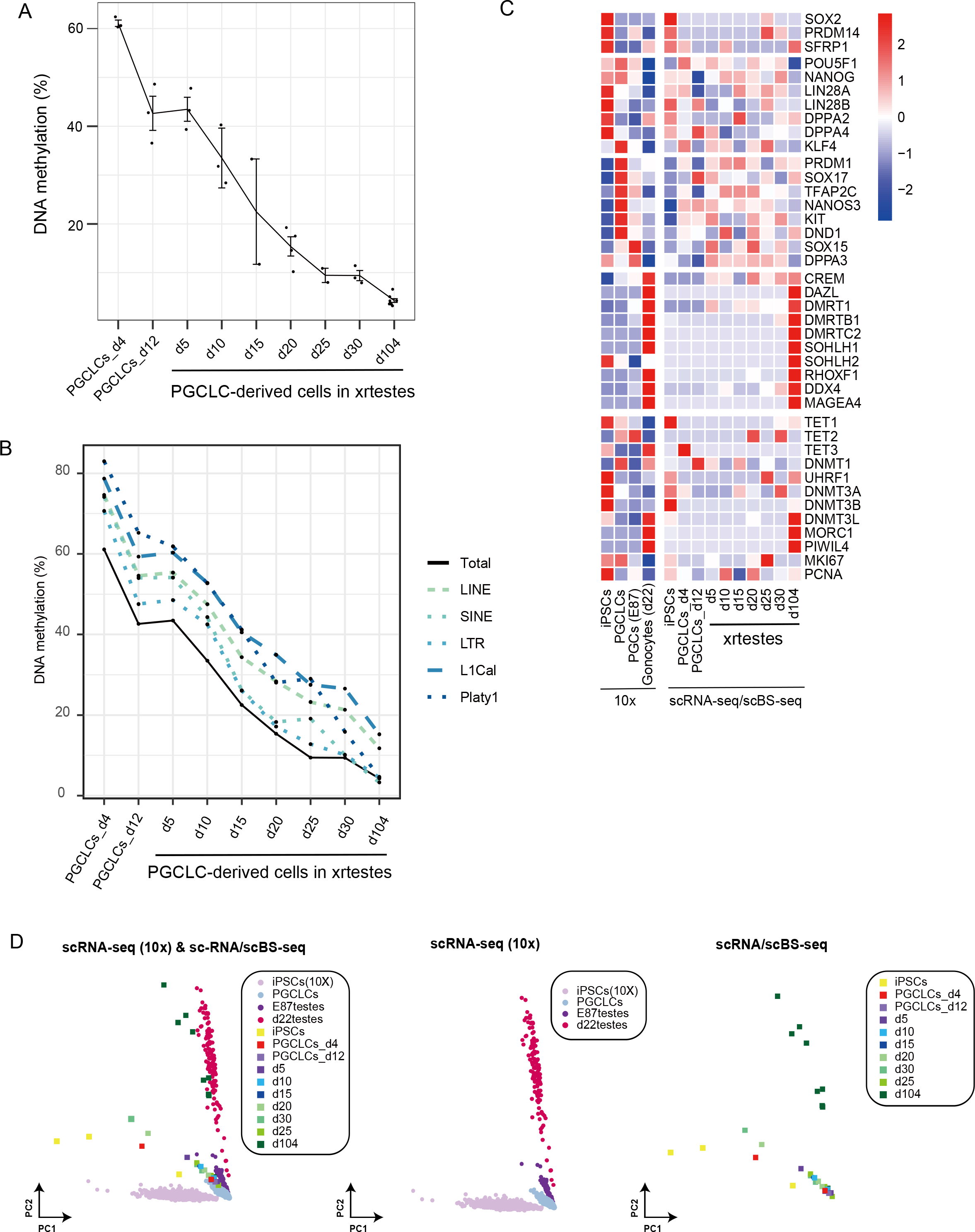
DNA methylation analyses during germ cell development from PGCLCs. **A** Single cell analysis of DNA methylation during PGCLC development. PGCLCs_d4 and PGCLCs_d12 correspond to 4-days and 12-days of floating aggregate culture respectively. d5-d104 represent duration (days) after transplantation into mouse kidneys. **B** Developmental dynamics of DNA methylation levels of retrotransposons. The average values are shown. **C** Heatmap analyses of marker genes. **D** The data obtained by 10x and scRNA-seq/scBS-seq are shown together (left) or separately (center and right). The anchoring function was not used to integrate datasets of different platforms. There is a small displacement between the iPSCs of the 10x platform and iPSCs of the scRNA-seq/scBS-seq platform.

DNA methylation plays a critical role in the repression of retrotransposons in germline cells. In both mice and humans, active and young retrotransposons (e.g., IAP and LINE1 in mice, Alu and LINE1 in humans) show relatively high levels of residual DNA methylation in demethylated PGC genomes (Guo et al., 2015; Kobayashi et al., 2013; Seisenberger et al., 2012). Two types of potentially active retrotransposons exist in the marmoset genome. One is the LINE1 element, and the other is a very short ~100-bp SINE element named Platy-1 (Konkel et al., 2016). The DNA methylation dynamics of these two active retrotransposons (LINE1 and Platy-1) and three major classes of retrotransposons sequences (LINE, LTR, and SINE) were examined. In d4_PGCLCs, all three major retrotransposon sequences (LINE, 76.6%; LTR, 70.6%; SINE, 74.1%) showed higher levels than the genome average (61.1%), and the two active retrotransposons (LINE1: 78.7%, Platy-1: 82.9 %) showed the highest levels. As PGCLC development progressed, all retrotransposons lost DNA methylation with dynamics similar to those of the genomic average. In d104_xrtestes, DNA methylation levels of all the examined retrotransposons were decreased more than five-fold (LINE1, 15.2% from 76.6%) to 17-fold (Platy-1, 4.7% from 82.9%). LINE1 (15.2%) and LINE (11.8%) still showed much higher levels than the genomic average (4.3%). On the other hands, other retrotransposons (LTR, 4.5%; SINE, 3.3%; Platy-1, 4.7%) showed similar levels to the genomic average. Thus, a higher level of residual methylation was retained in LINE1 but not in Platy-1.

To correlate germ cell development in xrtestes with *in vivo* germ cell development, we analyzed the RNA expression of PGCLC-derived germ cells, in which we analyzed DNA methylation. Upon differentiation of iPSCs into PGCLCs, the expression of *UHRF1* (involved in DNA methylation maintenance) and *de novo* DNA methyltransferase *DNMT3A/3B/3L* decreased (Figure 5C). This decrease may be involved in the demethylation of the PGCLC genome. In d104_xrtestes, *DNMT3L, PIWIL4*, and *MORC1* were highly upregulated, establishing the stage for *de novo* methylation of retrotransposons. In addition, the expression of gonocyte genes (*CREM, DAZL, DMRT1, DMRTB1, DMRTC2, SOHLH1, SOHLH2, RHOXF1, DDX4*, and *MAGEA4*) was observed in d104_xrtestes as well as in gonocytes from *in vivo* d22 testes (Figure 5C). Some of them (*CREM, DMRT1, DMRTB1*, and *DDX4*) are expressed from earlier stage of xrtestis development. These genes were also detected in *in vivo* PGCs from E87 testes (Figure 5C), suggesting that developmentally regulated gene expression is recapitulated in the xrtestis system. PCA revealed that PGCLC-derived cells, except those from d104_xrtestes, closely aligned with E87 testis germ cells (Figure 5D). On the other hand, PGCLC-derived germ cells from d104_xrtestes clustered together with 22d testis germ cells. PGCLCs differentiate into gonocyte-like cells in the xrtestis.

## Discussion

In this study, marmoset PGCLCs were generated from iPSCs using an mRNA-transfection-based method. This is the first report of PGCLC generation using mRNAs. Since this method is simple and efficient, it may also be useful for other species. Furthermore, the generated PGCLCs differentiated into gonocyte-like cells in the xrtestes that were transplanted under the kidney capsules of immunodeficient mice. Stepwise expression of PGC and gonocyte marker genes was observed. DNA methylation was progressively lost and almost completely erased in the gonocyte-like cells. Thus, early germ cell development *in vivo* was recapitulated by our PGCLC-initiated system. This study provides a platform for developmental studies on marmoset germ cells and the generation of genetically modified marmosets.

We induced PGCLCs from iPSCs using a combination of *SOX17* mRNA transfection and subsequent floating aggregate culture. Our method was based on a report on PGCLC generation by *SOX17* overexpression using an inducible system (Irie *et al*., 2015; Kobayashi *et al*., 2017), which requires prior transgene integration. To omit this step, we used mRNA transfection-based overexpression. Although the induction rate was highly dependent on the iPSC lines (data not shown), as in humans (Chen et al., 2017), our induction efficiency usually reached >80% when highly competent lines were used (Figure 1B, S1). Since this efficiency is comparable to or higher than those of existing methods (Irie *et al*., 2015; Jo et al., 2022; Kobayashi *et al*., 2017; Sakai *et al*., 2020; Sasaki *et al*., 2015; Sosa *et al*., 2018; Yoshimatsu *et al*., 2021), we believe that the mRNA transfection method reported here serves as an alternative method.

After developing a solid foundation for PGCLC induction, we aimed to differentiate PGCLCs into a more advanced state. Matoba et al. reported that *in vivo* mouse PGCs developed into spermatids in reconstituted testes transplanted under the kidney capsule (Matoba and Ogura, 2011). This led us to examine whether immunodeficient mouse kidneys serve as suitable sites to develop xrtestes. The xrtestes developed well under the kidney capsule. Using this technique, PGCLCs in the xrtestes were found to differentiate into gonocyte-like cells. All gonocyte-like cells from the d82_xrtestes were negative for the PGC marker TFAP2C (Figure 4). Given that half of the germ cells still express TFAP2C in newborn testes (Mitchell *et al*., 2008), gonocyte-like cells in d82_xrtestes are more developmentally advanced than newborn testis germ cells. However, our bisulfite-seq analyses showed that PGCLC-derived cells in d82_ and d104_xrtestes did not undergo *de novo* DNA methylation (Figure 5). In marmoset testes, *de novo* DNA methylation is initiated at 4 months old at the latest (Langenstroth-Rower et al., 2017), although the precise timing has not yet been determined. Therefore, gonocyte-like cells in d82_ and d104_xrtestes likely correspond to *in vivo* gonocytes in newborn and 4-month-old animals. The kidney transplantation method reported here will be a robust *in vivo* method for male PGCLC differentiation in other species as well. Reconstituted embryonic ovaries from mice and cynomolgus monkeys develop well in mouse kidneys (Matoba and Ogura, 2011; Mizuta et al., 2022). Therefore, the kidney transplantation of (xeno)reconstituted ovaries may be also useful for advancement of female PGCLC development in marmosets and other species.

Successful marmoset PGCLC induction has been previously reported by Yoshimatsu et al. (Yoshimatsu *et al*., 2021). However, their induction efficiency (~40% for the two ESC lines and 1–2% for the two iPSC lines) seemed to be not as high as ours. In addition, their study did not test the developmental potential of these PGCLCs. By contrast, PGCLCs were differentiated into gonocyte-like cells in our study. Their methods involved both transgene (*SOX17* and *BLIMP1*) overexpression and pre-ME/iMeLC induction steps, and it took 10 days (+prior transgene integration) for the procedure. Our method required only 6 days, and no prior transgene integration is required. Furthermore, we did not use *BLIMP1*, because the addition of *BLIMP1* mRNAs to *SOX17* mRNAs did not have any positive effect on PGCLC induction in our system (Figure S7). Combinatorial expression of *BLIMP1* and *SOX17* has been reported to promote PGCLC in humans (Kobayashi *et al*., 2017). Species difference or different methods used likely account for the discrepancy of the effect of *BLIMP1*. Recently, another group reported marmoset PGCLC induction from PSCs (Seita et al., 2022). Their induction efficiency was 40% at the highest using a similar method reported in Cynomolgus monkeys and rabbits (Kobayashi et al., 2021; Sakai *et al*., 2020). They cultured PSCs in the presence of the WNT inhibitor IWR-1, and PGCLCs were directly induced from PSCs without undergoing Pre-ME/iMeLC. Although their PGCLCs differentiated into DDX4-positive cells (corresponding to late PGCs or gonocytes), complete DNA demethylation and the potential for differentiation into MAGEA4-positive gonocyte-like cells were not examined. Thus, our study provides two efficient and useful systems associated with marmoset PGCLCs: (1) an mRNA-transfection-based PGCLC induction system and (2) a kidney transplantation-based PGCLC to gonocyte-like cell differentiation system.

The gonocyte-like cells generated in this study require further development for the generation of functional gametes. The next step is further differentiation into late gonocyte-like cells that undergo *de novo* DNA methylation. It is important to understand the cues that initiate *de novo* DNA methylation. Furthermore, the current protocol requires a long time to differentiate gonocyte-like cells from PGCLCs. Shortening this time is also an important next step. However, undertaking the normal demethylation process in PGCs is likely important for generating functional gametes. In fact, bypassing this resulted in abnormal DNA methylation patterns in mouse oocytes (Hamazaki et al., 2021). Furthermore, *in vitro* PGCLC culture and the resultant prior erasure of DNA methylation have been reported to be essential for the spermatogenic potential of PGCLC-derived spermatogonial stem cells (Ishikura et al., 2021). DNA methylation dynamics revealed in this study are, in part, useful for determining which developmental stage can be bypassed without affecting the DNA demethylation process. Our study provides a solid foundation for complete generation of gametes from pluripotent stem cells.

### Marmoset housing and samplings

All animal experiments using marmosets and mice were approved by the Animal Committee of the Central Institute for Experimental Animals (Approval number; 17029A, 18031A, 19033A, 21002A, 21012A, and 21052A). Marmosets (CLEA Japan) were housed in Central Institute for Experimental Animals. The marmosets were housed in stainless steel cages (W436 × D750 × H765 mm to W910 × D750 × H2050 mm) under the following conditions: temperature, 27 °C, humidity 40%, room pressure, +20 hPa, light 12 h per day, and basic food CMS-1 (CLEA Japan).

To obtain fetal gonads (E74 and E82 ovaries and E87 testes), frozen or fresh early embryos (8-cell to morula) were transferred into recipient uteri (Takahashi et al., 2014). The developmental days were determined based on the day of ovulation when the serum progesterone of the recipient animals exceeded 10 ng/ml. Fetuses were obtained by C-sections. The pregnant female marmosets were pre-anesthetized with 0.04 mg/kg medetomidine (Domitor, Nippon Zenyaku Kogyo), 0.40 mg/kg midazolam (Dormicam, Astellas Pharma), and 0.40 mg/kg butorphanol (Vetorphale, Meiji Seika Pharma), then sedated by isoflurane (Isoflurane inhalation solution, Viatris) inhalation. While C-section, the animals were warmed on a warmer pad on the surgery table. The uterus was exteriorized via a midline laparotomy and lifted from the abdominal cavity, and the fetus was removed from the uterus. After C-section, the female marmoset’s abdominal wall and skin were sutured and kept warm in the intensive care unit until awakened from anesthesia.

Testis from 22d marmosets was obtained by hemicastration under anesthesia by inhalation of 1–3% isoflurane. Newborn ovaries and adult testes were obtained from animals sacrificed for use in other experiments and for illnesses that could not be cured, respectively. The list of animals and cell lines used in this study is shown in Table S1.

### Cell culture

Marmoset iPSCs were cultured in MEF-conditioned primate ES cell medium (RCHEMD001; REPROCELL) containing bFGF (RCHEOT003; REPROCELL) and 1× antibiotic-antimycotic (0289-54; Nacalai Tesque)(Watanabe *et al*., 2019). Thereafter, the iPSCs were dissociated into single cells and passaged by treatment using Accumax (17087-54; Nacalai Tesque) at 37 °C and then plated 5.0 × 10^4^ cells in one well of 6-well cell culture plates with conditioned medium containing iMatrix (892021; FUJIFILM) and 10 μM Y-27632 (08945-42; Nacalai Tesque).

### Induction of PGCLCs

iPSCs (lines 971 and 972) were induced from ear fibroblast cells as described previously (Watanabe *et al*., 2019). Lipofection-based mRNA transfection was performed to generate PGCLCs from iPSCs. The day before transfection (day 0), 5.0 × 10^4^ marmoset iPSCs were plated in 12-well cell culture plates. The next day (day 1), the medium was replaced with 500 μL of the new conditioned medium. For transfection, two tubes containing 62.5 μL of OPTI-MEM (A4124801; Invitrogen) were prepared. In one tube, RNAs [0.2 μg of SOX17 mRNA, 0.05 μg of P53DD mRNA, 0.075 μg of the mixture of E3, K3, B18R mRNAs (00-0076; Reprocell)] were added. In the other tube, 1.5 μL of Lipofectoamine RNAiMax Transfection Reagent (13778150; Invitrogen) was added. After mixing well, the solutions in the two tubes were mixed, and the mixture was allowed to stand for 10 min. Next, the mixture was added dropwise into one well of a 12-well plate. The next day (day 2), the medium was replaced with 500 μL of new conditioned medium, and RNA transfection was performed again, after which the medium was changed, and RNA transfection was performed at 10 am and 4 pm.

On the morning of day 3 (~10 am), cells were dissociated using Accumax. Subsequently, ~5,000 cells were plated in each well of a Nunclon Spher 96-Well plate (174929; Thermo Fisher Scientific) in 100 μL of aRB27 (1% B27/2 mM L-glutamine/1× non-essential amino acids/1× antibiotic-antimycotic in advanced RPMI medium) containing 50 ng/mL EGF (236-EG-200; R&D Systems), 100 ng/mL SCF (455-MC-050; R&D Systems), 1,000 U/mL LIF (LIF1010; Merck), 400 ng/mL BMP4 (314-BP-500; R&D Systems), and 10 μM Y-27632. The medium was changed every 4 days.

### sc library generation and data analyses

The iPS cells were dissociated using Accumax, while PGCLCs, ovaries (E74, E82, and newborn), and testes (E87 and 22d) were dissociated using 0.25% trypsin EDTA, and adult testes (3 years 10 months) were digested by stepwise treatment with collagenase 1 (1mg/mL) and 0.25% trypsin EDTA. For iPS cells, PGCLCs, and E82 ovaries, libraries were constructed using a Chromium Next GEM Single Cell 3′ GEM Library & Gel Bead Kit v3 (10x Genomics). Other libraries were constructed using a Chromium Single Cell 3′ Library and Gel Bead Kit v2 (10x Genomics), and sequencing was performed using a HiSeq4000 system (Illumina). The sample information is summarized in Table S1. See supplementary information for the scRNA-seq and scBS-seq library generation.

To analyze s 10x data, datasets were mapped to the common marmoset genome, calJac4 (calJac3 for Fig. 2C and D), using CellRanger. For annotation of the calJac4 genome, NCBI Callithrix jacchus Annotation Release 105 was used, and UMI count data were analyzed using Seurat. Normalization and standardization were performed using *NormalizeData* and *ScaleData* functions. Principal component analysis was performed using the inbuilt *RunPCA* function. Dimensionality reduction was visualized in 2D using *DimPlot*. See supplementary information for the details on Fig. 2C, D, 5C, and D.

10x single cell data were deposited to ArrayExpress (Accession No. E-MTAB-12123) and scRNA-seq/scBS-seq data were registered to DDBJ (Accession No. DRA014666 and DRA014672).

### Karyotype analysis

The iPSCs were cultured for 3 h in a medium containing 0.1 μg/mL colcemid (15212012; Gibco) and dissociated into single cells by treatment with Accumax. After collection, the cells were treated with 0.075 M KCl for 30 min at 25°C. Subsequently, the cells were fixed using a fixing solution (70% methanol/30% acetic acid). The fixed cells were then spread onto a glass slide using HANABI and stained with Hoechst 33342 (H1399; Invitrogen). Karyotype images were obtained using a Leica 6000B microscope.

### *In vivo* differentiation

To isolate fetal testicular somatic cells, pregnant ICR females were sacrificed, and embryos were collected at E13.5. Fetal male testes were distinguished from ovaries by their appearance, and the mesonephros attached to the testes were removed using a tungsten needle. Thereafter, the isolated testes were treated with 0.25% trypsin EDTA (25200056; Nacalai Tesque) for 10 min at 37] for dissociation into single cells. To remove endogenous mouse PGCs, cells were plated in a cell culture plate in MEM α (12571063; Gibco) containing 10% KSR (10828-028; Gibco) and cultured for 6 h (Ohinata et al., 2009). PGCs usually do not attach to plates; the attached cells were removed from the culture plate using 0.25% trypsin/EDTA and collected as fetal testicular somatic cells.

Marmoset d4_PGCLCs were purified using *SOX17*-T2A-tdTomato fluorescence and FACS. The collected d4_PGCLCs (5.0 × 10^3^ cells) were mixed with E13.5 fetal testicular somatic cells (5.0 × 10^4^ cells) and plated in an ultra-low attachment 96-well plate (174929; Thermo Fisher Scientific) with MEM α containing 10% KSR, 1× antibiotic-antimycotic and 10 μM Y27632. The next day, cell aggregates were transplanted under the kidney capsule of NOG (NOD/Shi-scid, IL-2RγKO) mice to promote differentiation (Ito et al., 2002). Thereafter, NOG mice were anesthetized using medetomidine, midazolam, and butorphanol, and an incision was made in the peritoneum to expose the kidney capsule. Next, the kidney capsule was carefully incised using an injection needle under an inverted microscope, and cell aggregates were picked up using a glass capillary and transplanted into the kidney capsule.

### Immunofluorescence

Xrtestes were collected from kidneys, fixed in 4% PFA (09154-56; Nacalai Tesque), and then embedded into paraffin blocks. Afterward, xrtestes sections (4 μm) were deparaffinized and rehydrated using a xylene and ethanol series. For antigen retrieval, slides were heated to 95 °C for 10 min in 0.01 M citrate buffer (pH 6.0). The sections were blocked using a primary antibody diluent (AR9352; Leica Biosystems) and incubated overnight at 4 °C with the following primary antibodies: anti-GFP (308SS; 1:1,000; Novus Biologicals), anti-GFP (ab5450; 1:3,000; Abcam), anti-LAMININ (L9393; 1:200; Sigma-Aldrich), anti- 3β-HSD (sc-515120; 1:1,500; Santa Cruz Biotechnology Inc), anti-WT1 (ab89901; 1:500; Abcam), anti-MKI67 (NCL-ki67p; 1:50; Leica Biosystems), anti-TFAP2C (sc-12762; 1:100; Santa Cruz Biotechnology Inc), anti-DDX4 (AF2030; 1:500; R&D Systems), anti-MAGEA3/4 (MABC1150; 1:250; Merck), and anti-PIWIL4 (GP1831 produced in a guinea pig; AHSSFRATEVGRTQD-Cys; 1:200). Subsequently, the sections were incubated with species-specific secondary antibodies for 60 min at room temperature.

### RT-qPCR

Purified RNA was extracted from undifferentiated marmoset iPSCs, sorted d4_PGCLCs, and xrtestes using TRIzol Reagent (15596026; Invitrogen). RNA was converted to cDNA using Superscript IV Reverse Transcriptase (11766050; Invitrogen), and gene expression analysis was performed using the primers listed in Table S2.

## Acknowledgments

We thank Akihiro Umezawa and Hidenobu Soejima for their encouragement and Yasufumi Sakakibara and Kengo Sato for their support with computational resources and Fuchou Tang, Rui Wang, and Shin-ichi Tomizawa for single cell bisulfite-seq protocols and Akiyumi Tashiro for making comments on our manuscript and Kenji Kawai for making paraffin blocks and Haruka Shinohara for advice on karyotyping and veterinarians and animal technicians in CIEA for marmoset and mouse housing. This research was supported by AMED and KAKENHI under the grant numbers JP19gm6310010, JP20gm6310010, JP21gm6310010, JP22gm6310010 (AMED T.W.), JP18dm020765 (AMED E.S.), 20H05764, 20H03177, and 22K18356 (KAKENHI T.W.).

## Author contributions

M.I., C.D., and TW performed the experiments. C.P., T.B., M.I., and T.W. performed informatics analyses. M.I. and T.W. conceived of the study, designed the experiments, and wrote the manuscript. K.K., K.S., Y.T., and K.W. provided materials and information. E.S. shared the equipment and samples. H.Y., H.K., A.M., KK, M.I., and T.W. generated sequencing libraries. All authors read and approved the final manuscripts.

