## Supplementary file for "mRNA-based generation of marmoset PGCLCs capable of differentiation into gonocyte-like cells"

#### Table of contents

Supplemental Methods (pp. 2-6)

Legends for Supplemental Figures 1-7 (p. 7)

Supplemental References (p. 8)

Supplemental Figures 1-7 (pp. 9-15)

Supplemental Tables 1-2 (pp. 16-19)

**Supplemental Methods**

**mRNA preparation**

For *in vitro* transcription of *SOX17* mRNA, marmoset *SOX17* cDNA was PCR-amplified from testis cDNAs using a following set of primers: Sox17_5UTR and Sox17_R_Asc1. The PCR product was again amplified to include the human *HBA* gene 5' UTR by using a following set of primers: 5UTR_HBA_Not1 and Sox17_R_Asc1. The primer sequences are listed in Table S2. The product was digested using Asc1 and Not1 and then ligated to phBG vector (deposited in RIKEN BRC #RDB18330) to generate phBG-SOX17 vector. phBG-SOX17 and previously generated phBG-P53DD (Addgene #149707) vectors were digested with HindIII for *in vitro* transcription using mMESSAGE mMACHINE T7 Ultra Kit (Ambion). *In vitro* transcribed mRNAs were purified using LiCl precipitation, and they were used for the transfection.

**Lentivirus preparation and infection**

Lentivirus was prepared using a protocol on the RIKEN website (https://dnaconda.riken.jp/Form_PDF/lntPrepen.pdf). Briefly, CS-CA-GFP (RDB05964), pCMV-VSV-G-RSV-Rev (RDB04393), and pCAG-HIVgp (RDB04394) were co-transfected into HEK293T cells. The supernatant was collected to enrich the lentivirus using LentiX-concentrator (Takara, 631231). The virus was stored at -80$^{o}$C until use. For the infection of the virus, the virus was added to the medium together with 4 μg/ml of polybrene. Then, spinfection was performed at 800 xg for 30 minutes at 32$^{o}$C.

**Knock-in of reporter gene into SOX17 gene locus**

To visualize the generation of PGCLCs, T2A-tdTomato was inserted into the C-terminus of the *SOX17* gene. To make the targeting vector, 820-bp 5' and 669-bp 3' arms were amplified from the genome and inserted into the BamHI-digested KW1521 vector using an In-fusion HD cloning kit (Takara, 639648). The KW1521 vector contains T2A-tdTomato cassette and human beta-actin promoter driven Neomycin resistant gene cassette for the selection. For generating sgRNA-expressing vectors, PCR was performed using a common primer (sgRNA-Universal-rev-primer) and specific primers (pHL_Sox17_1, pHL_Sox17_2, pHL_Sox17_3). PCR products were then digested with EcoRI and BamHI, and ligated to the EcoRI and BamHI sites of pHL-H1-ccdB-mEF1a-RiH vector (Addgene #60601).

For homologous recombination, iPS cells were plated at the concentration of $8 \times 10^4$ cells per one well of 6-well plate at one day before the transfection (Day 0). Transfection was performed using the following plasmid DNAs [1 μg of targeting vector, 0.5 μg of pHL-EF1a-SphcCas9-iP-A (Addgene #60599), and 167 ng each of three sgRNA vectors (pHL_H1_Sox17_1, pHL_H1_Sox17_2, pHL_H1_Sox17_3)] using Lipofectoamine 3000 reagent (Thermo Fisher Scientific, L3000008) or Lipofectoamine stem transfection reagent (Thermo Fisher Scientific, STEM00015) according to the manufacture's protocol. The day after the transfection (Day 2), cells were newly plated onto one well of a six-well plate. From Day 3, the selection was performed using 150 µg/ml G418. Colony pickup was performed ~Day 12.

For isolation of DNA for genotyping, cells were treated with DNA isolation buffer [100 mM Tris-HCl (PH 7.6)/100 mM NaCl/10 mM EDTA (PH 8.0)/0.5% SDS] containing 100 µg/ml Proteinase K at 50°C. After the phenol-chloroform extraction, DNA was precipitated by adding an equal volume of isopropanol at room temperature. PCR was performed to detect knock-in using the following sets of primers: Sox17_scr5F and tdTomatoR1 (recombination at the coding region side), Sox17_scr3R and human-b-actProR (recombination at the 3' UTR side), and Sox17_scr5F and Sox17_scr3R (for detecting entire knock-in insertion or wildtype bands).

**FACS purification of PGCLCs**

PGCLC aggregates were dissociated with 0.25% Trypsin-EDTA for 10 minutes at 37 °C. The reaction was stopped by adding an equal amount of 10% KSR/alpha-MEM. DNaseI was added at the final concentration of 25 µg/ml, and cells were left at room temperature for 1 minute before centrifuge. After adding FACS buffer containing DAPI for removing dead cells, cells were filtered using a 60 µm cell strainer. tdTomato positive cells were purified using SONY SH800 using purity mode.

**Comparison of *in vitro* datasets with in vivo dataset and Human datasets (Figure 2D, E)**

Marmoset 10X datasets were mapped to the common marmoset genome (Callithrix jacchus 3.2.1) using CellRanger 3.0.2, which was used filter out low-quality cells and empty droplets. Gene annotation files for common marmoset were downloaded from Ensembl (release 91) Gene annotation files for common marmoset genome (Caljac 3.2.1) were downloaded from Ensembl (release 91) and gene models extended similarly to the approach that was reported[1,2].

For comparison with human, several existing human 10x datasets were also analyzed, including germ cell samples from neonatal testes[3] (GSE124263) and PGCLCs induced from iPSCs[4] (GSE140021). Smart-seq2 datasets of male and female fetal germ cells[5] (GSE86146) were incorporated in downstream analyses alongside single cell data from human iPSC and PGCLCs[6] (GSE99350).

All samples that passed QC were analyzed using Seurat v3.1.2[7]. UMI data was normalized and standardised using the *NormalizeData* and *ScaleData* function using either the top 2,000, 5000 or 20,000 most varied genes for initial analysis. Principal component analysis was run using the inbuilt *RunPCA* function, with nonlinear dimensionality reduction techniques generated using *RunUMAP*. Dimensionality reduction was visualised in 2D using *DimPlot*.

Data from marmoset and humans were incorporated together for an integrative view of germ cell development. First our 10X samples of marmoset germ cells were integrated with iPSCs and PGLCs to create a marmoset reference dataset. 10X samples from human neonate gem cell lineages were merged with 10X samples of human iPSCs and PGCLCs. Smart-Seq2 datasets from male and female germ cells were merged with SS2 samples of PGCLCs and iPSCs. Each gene in the marmoset data was annotated by its human orthologue based on the Ensembl BioMart database (https://m.ensembl.org/info/data/biomart/index.html). These homologous genes were used for the integrative analyses of Human and marmoset data. The three datasets were then jointly aligned in Seurat based on Canonical Correlation Analysis (CCA) and mutual nearest neighbor (MNN) approaches. Specifically, *FindIntegrationAnchors* was run using 4000 features and *IntegrateData* (with 20 dimensions) was used to calculate corrected gene expression matrices for the three datasets. Datasets were visualized using PCA on the corrected gene expression matrix.

Cell lineages for marmosets were first identified in individual samples based on the expression of known marker genes (Figure S2). Unbiased clustering on the aligned marmoset-human dataset was done using the *FindClusters* function, and individual clusters were annotated based on the dominant lineages in the cluster. These annotations were additionally checked against the aligned human annotations and showed strong agreement.

Finally, marmoset lineages were appended by developmental stage, and the average expression of lineages was calculated using *AverageExpression*. Row normalized expression values for an extended panel of known markers was visualized using pheatmap.

**RNA and DNA purification from the same single cell**

xrtestes were dissociated using 0.25% Trypisn-EDTA. xrtestis cells were resuspended in 0.04% BSA in PBS for single-cell pick up. Single-cell was picked up by using a mouth pipet and put into a PCR tube. Lysis buffer (1.14 U/μl Rnase Inhibitor, 0.54% Triton X-100) containing DynaBeads MyOne Carboxylic acid (Thermo Fisher) was added. Beads bound to the nucleus and supernatant corresponding to the cytoplasmic fraction were separated. DNA isolation buffer (20 mM Tris-EDTA, 20 mM KCl, 0.3% Triton X-100, 1mg/ml Proteinase K, 0.1 pg/μl λDNA, 2 ng/μl carrier RNA) was added to the beads, and they were incubated at 50°C for 10 minutes. Proteinase was then heat-inactivated by 75°C for 30 minutes.

**Single-cell RNA-seq of the cytoplasmic fraction**

Before cDNA synthesis, ERCC RNA (Invitrogen, 1:500,000) was added to the cytoplasmic RNA. cDNA was made from cytoplasmic RNAs using oligo dT primer with cell barcode and UMI sequences (CB16UMI12-RT), template switch (TS) oligo DNA, and SuperScript II Reverse transcriptase (Thermo Fisher). To prevent concatamerization of the TS oligo, non-natural nucleotides were added to its 5' end. PCR reaction was performed using 3' Anchoerd primer and ISPCR primers to amplify cDNA. After purification of cDNA using AMPure beads, the qPCR reaction was carried out using a primer pair that detected germ cell-specific gene *DDX4* for the selection of germ cell. After pooling 8 cDNAs with different barcodes, PCR was performed using a biotinylated primer bound to 5' end of oligodT sequence (Biotin-index-primer) and unmodified primer annealed to 5' end of TS oligo (ISPCR). Fragmentation was carried out using NEBNext Ultra II FS DNA Library Prep Kit for Illumina, and biotinylated DNA was purified using streptavidin beads. Adapter ligation and PCR-amplification (using QP2 and Modified_P5_primer) were then performed according to NEBNext Ultra II FS DNA Library Prep Kit for Illumina. For PCR amplification After quality check using Bioanalyzer (Agilent), paired-end 150-bp sequencing was performed by Illumina HiSeqX. Primer sequences are listed in Table S2.

**Single-cell BS-seq**

Single-cell BS-seq was conducted based on published literatures[8-10]. Before the bisulfite reaction, non-methylated lamda DNA (Promega) was added to the DNA. For bisulfite conversion, MethylCode Bisulfite Conversion kit (Invitrogen) was used. For making single-cell BS-seq libraries, two rounds of random priming reaction were performed using a random primer with a nucleotide ratio of A:T:G:C = 4:4:1:1 (scBS-P5-N9-oligo1), which reduces bias toward preferential amplification of GC-rich sequences. After ExonucleaseI treatment for removing the remaining random primer, second-strand synthesis was performed using scBS-oligo2 primer. PCR amplification were then carried out using unique dual index primers (NEBNext Multiplex Oligos for Illumina, 96 Unique Dual Index). After quality check using a bioanalyzer (Agilent), paired-end 150-bp sequencing was performed by Illumina HiseqX.

**scRNA/scBS-seq data analysis**

Our custom scRNA libraries were designed for analyses using CellRanger. Prior to the analyses using CellRanger, Read1 and Read2 were switched by changing the file names. Datasets were mapped to the common marmoset genome calJac4 using CellRanger. For annotation of calJac4 genome, NCBI Callithrix jacchus Annotation Release 105 was used. UMI count data were analyzed using Seurat. The count matrix was created by *CreateSeuratObject* function. To extract the cells of interest, a *subset* function was used. Data from different libraries were combined using the *merge* function. To remove cells with a small number of UMIs, a subset function was used by setting a cutoff value of unique feature counts over 1500. Normalization and standardization were performed using the *NormalizeData* and *ScaleData* functions, respectively. Appropriate cell names were reassigned using the *changeindent* function. Variable genes were extracted by *FindVariableFeatures* and principal component analysis was run using the inbuilt *RunPCA* function. Heatmap was generated using a *pheatmap* package.

Quality of the library was checked by FastQC and trimmed away adaptor sequence using the TrimGalore program using an option --quality 20 --stringency 3 --length 50 --clip_R1 9 --clip_R2 9 --paired --trim1 --phred33. Reads were mapped to the reference genome using Bismark with a command bismark --bowtie2 --fastq --non_directional --un. Duplicate reads were removed with *deduplicate* function in Bismark (deduplicate_bismark –bam). Deduplicated BAM files were then converted to bedGraph using *methylation extractor* (bismark_methylation_extractor -p --bedGraph -counts).

For retrotransposon analyses, the coordinates of 8,251 full-length LINE1 and 2,275 full length Platy-1 elements were retrieved from L1 base2 (l1base.charite.de/l1base.php) and a published literature[11], respectively. The calJac3 coordinates were converted to calJac4 coordinates using a liftover tool in UCSC (Lift Genome Annotations (ucsc.edu)). The coordinates for three major retrotransposon classes were retrieved from repeatmasker file downloaded from UCSC. Methylation data in bedgraph file format were used to calculate the average methylation levels.

#### Supplemental Figure Legends

**Figure S1. PGCLC generation from 971-STCE and 972-STCE iPSCs**

(A) Sox17-tdTomato (ST) and CAG-EGFP (CE) expression in d4_PGCLCs from 971-STCE and 972-STCE iPS cells. Scale bars: 100 μm (top) and 50 μm (bottom). (B) FACS analysis of 4d_PGCLCs from 971-STCE and 972-STCE iPS cells. Many cells are positive for Sox17-tdTomato fluorescence. Antibodies for BV421 (Y-axis) fluorescence were not used here.

**Figure S2. Determining the conditions for PGCLC induction using mRNAs**

**A**. Number of transfections. In each well of 12-well plate, $8x10^4$ iPSCs (mRNA) were seeded. After the indicated number of mRNA transfections, cells were plated onto the low binding 96 well plate. Error bars represent S.E. (N=2 or 3). **B.** Cell number of iPSCs (mRNA-ST) used for transfection. PGCLC induction was performed using a standard procedure (two successive day mRNA transfection and aggregate culture for four days).

**Figure S3. Single cell RNA-seq analyses of developing marmoset ovaries (A-C) and testes (D-F)**

Germ cell populations used for PGCLC developmental analyses are indicated.

**Figure S4. The expression of genes involved in sex differentiation in E74 ovaries and testes**

**Figure S5. The expression of PGC and gonocyte marker genes in xrtestes**

Immunofluorecent analyses of marker gene expression in developing xrtestes (PGCLCs from 971-STCE iPSCs). Scale bar: 20 um. PIWIL4-positive cells are indicated by white arrows.

**Figure S6. qPCR analyses of PGC and gonocyte marker genes in xrtestes**

qPCR analyses of marker gene expression in developing xrtestes (971-STCE and 972-STCE). Only one sample was examined in each stage, and the indicated values are the average of technical duplicates (N=2). Error bars represent S.D. The expression level was normalized to that of *GAPDH*. The value was further normalized to the expression level in the marmoset adult testes. The expression level in the adult testes corresponds to 1.

**Figure S7. Addition of BLIMP1 mRNA to SOX17 mRNA does not promote PGCLC induction**

*SOX17* and *BLIMP1* mRNAs were together transfected at various ratios. After the two successive day transfection, PGCLCs were generated using floating aggregate culture. d4_PGCLCs were used for the analyses. Error bars represent S.E. (N=3).

Figure S1

A

971-STCE（PGCLCs day4)

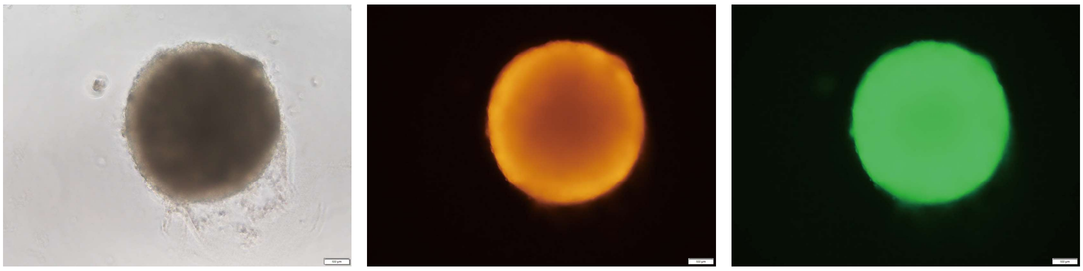

972-STCE（PGCLCs day4)

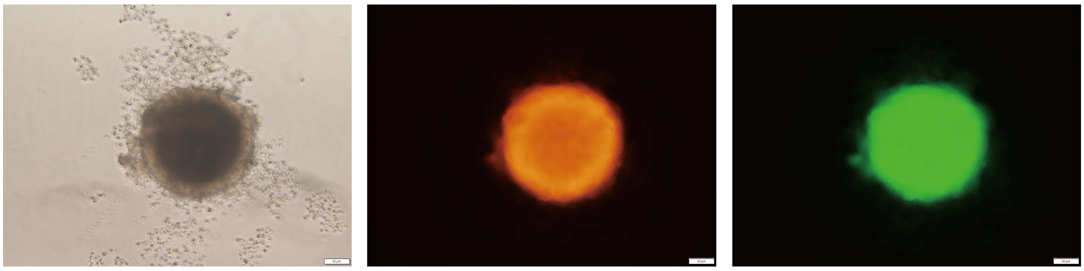

B

971-STCE FACS

972-STCE FACS

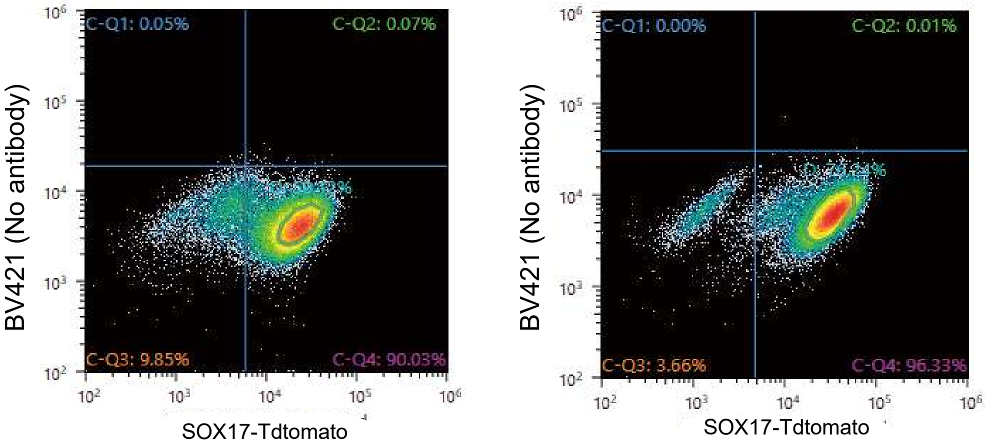

Figure S2

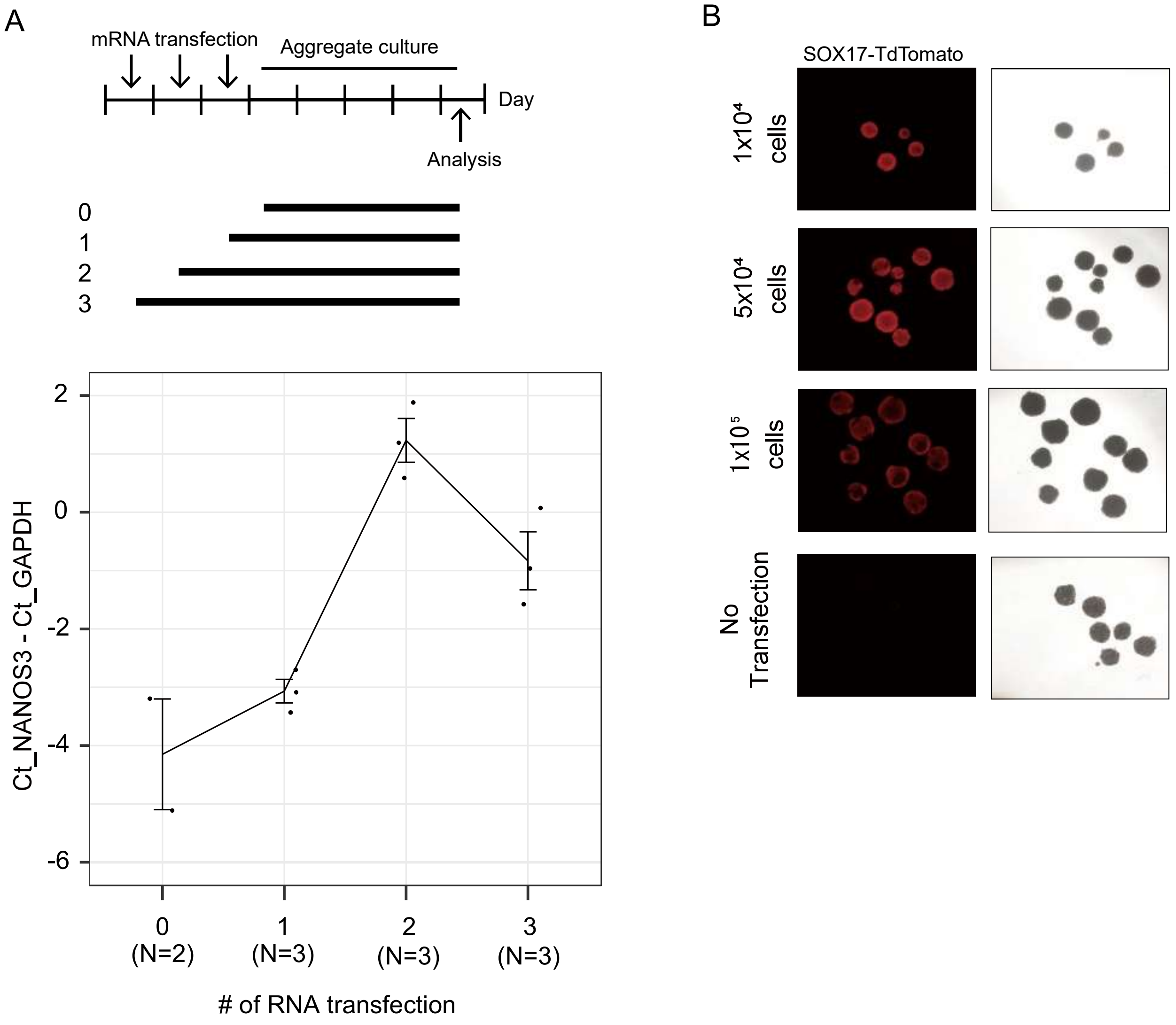
A
mRNA transfection
Aggregate culture
Day
Analysis
0
1
2
3
Ct_NANOS3 - Ct_GAPDH
2
0
-2
-4
-6
0
(N=2)
1
(N=3)
2
(N=3)
3
(N=3)
### of RNA transfection
B
SOX17-TdTomato
1x10⁴ cells
5x10⁴ cells
1x10⁵ cells
No Transfection

Figure S3

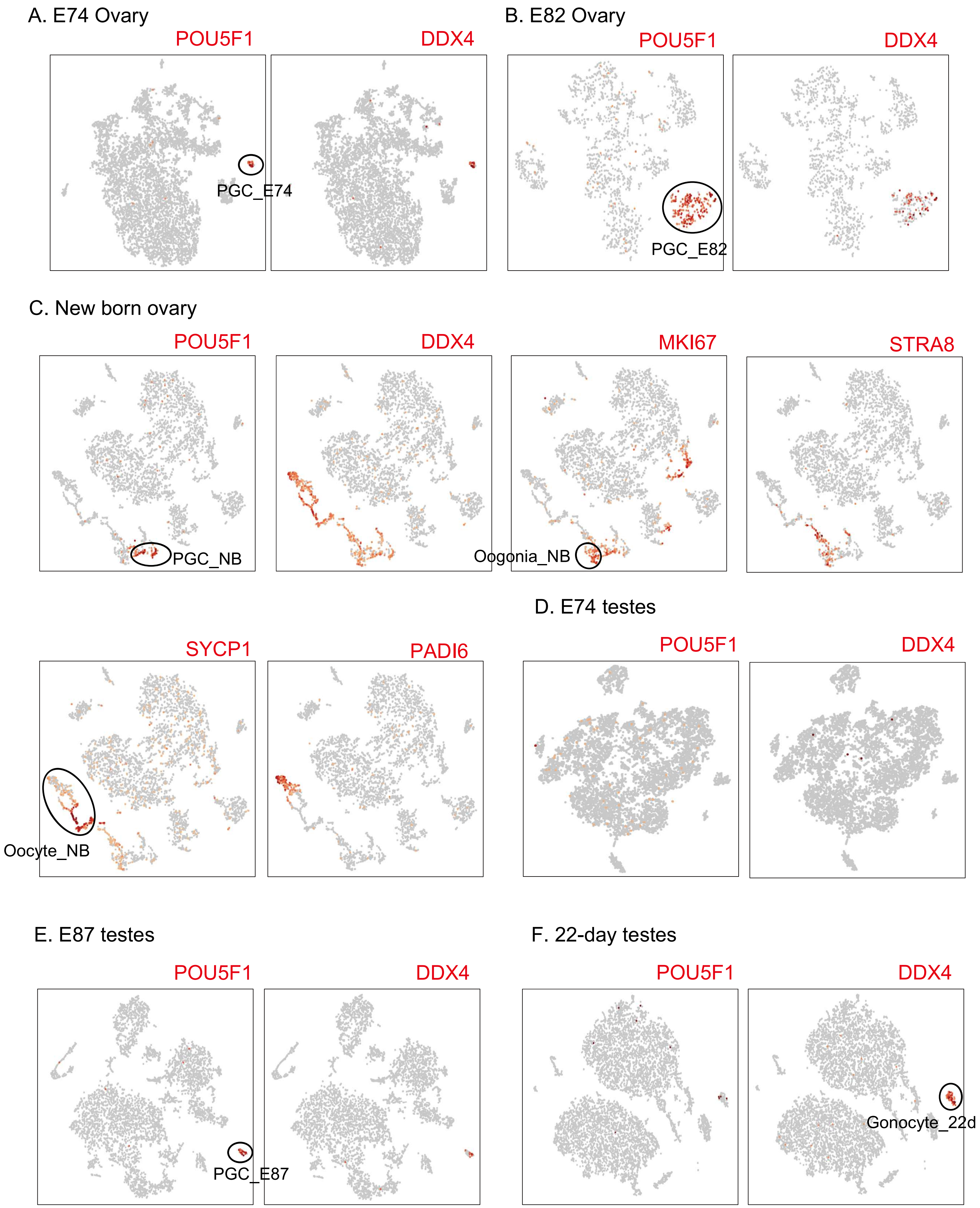
A. E74 Ovary
POU5F1
DDX4
PGC_E74
B. E82 Ovary
POU5F1
DDX4
PGC_E82
C. New born ovary
POU5F1
DDX4
MKI67
STRA8
PGC_NB
Oogonia_NB
SYCP1
PADI6
Oocyte_NB
D. E74 testes
POU5F1
DDX4
E. E87 testes
POU5F1
DDX4
PGC_E87
F. 22-day testes
POU5F1
DDX4
Gonocyte_22d

Figure S4

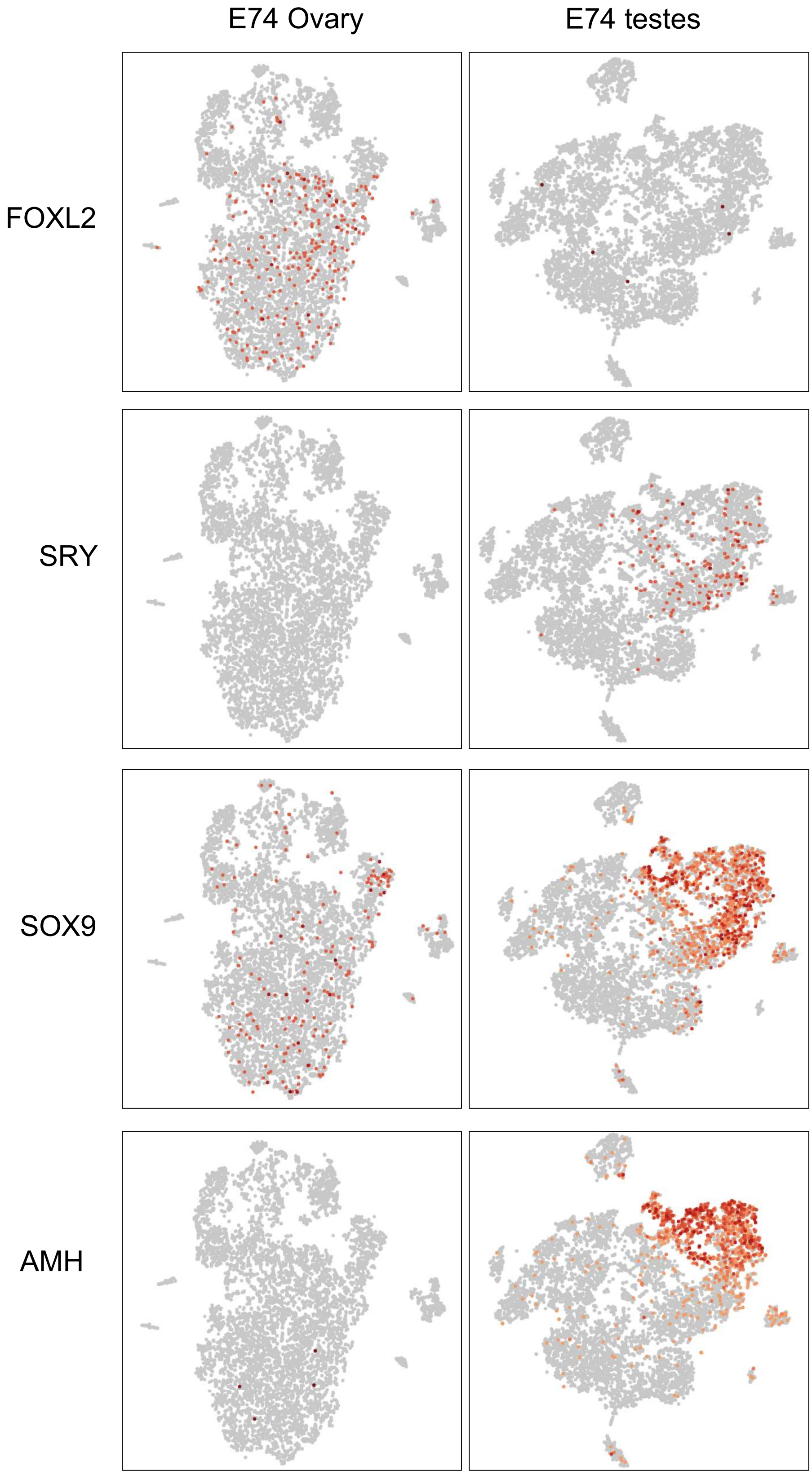

Figure S5

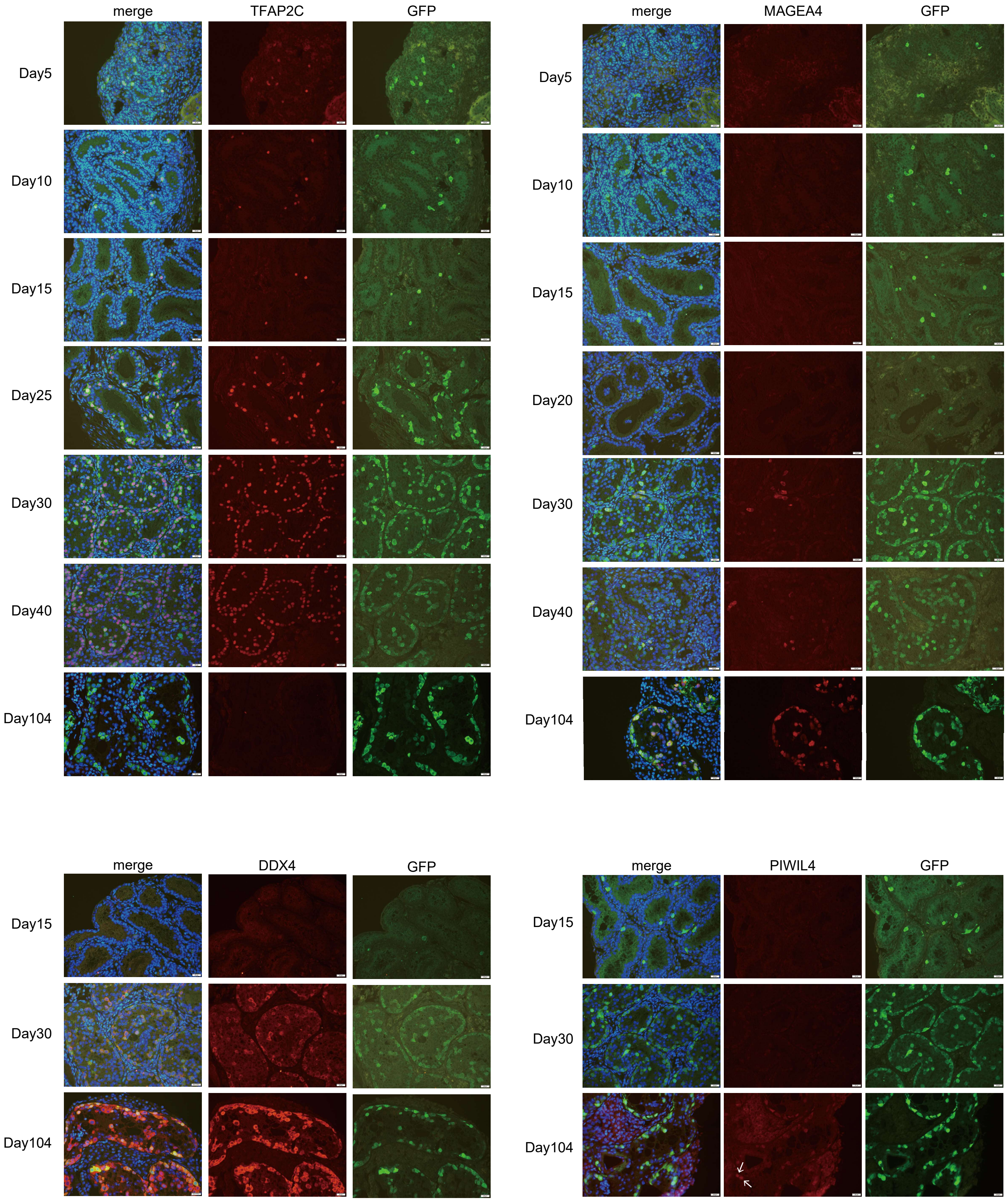
merge
TFAP2C
GFP
Day5
Day10
Day15
Day25
Day30
Day40
Day104
merge
MAGEA4
GFP
Day5
Day10
Day15
Day20
Day30
Day40
Day104
merge
DDX4
GFP
Day15
Day30
Day104
merge
PIWIL4
GFP
Day15
Day30
Day104

Figure S6

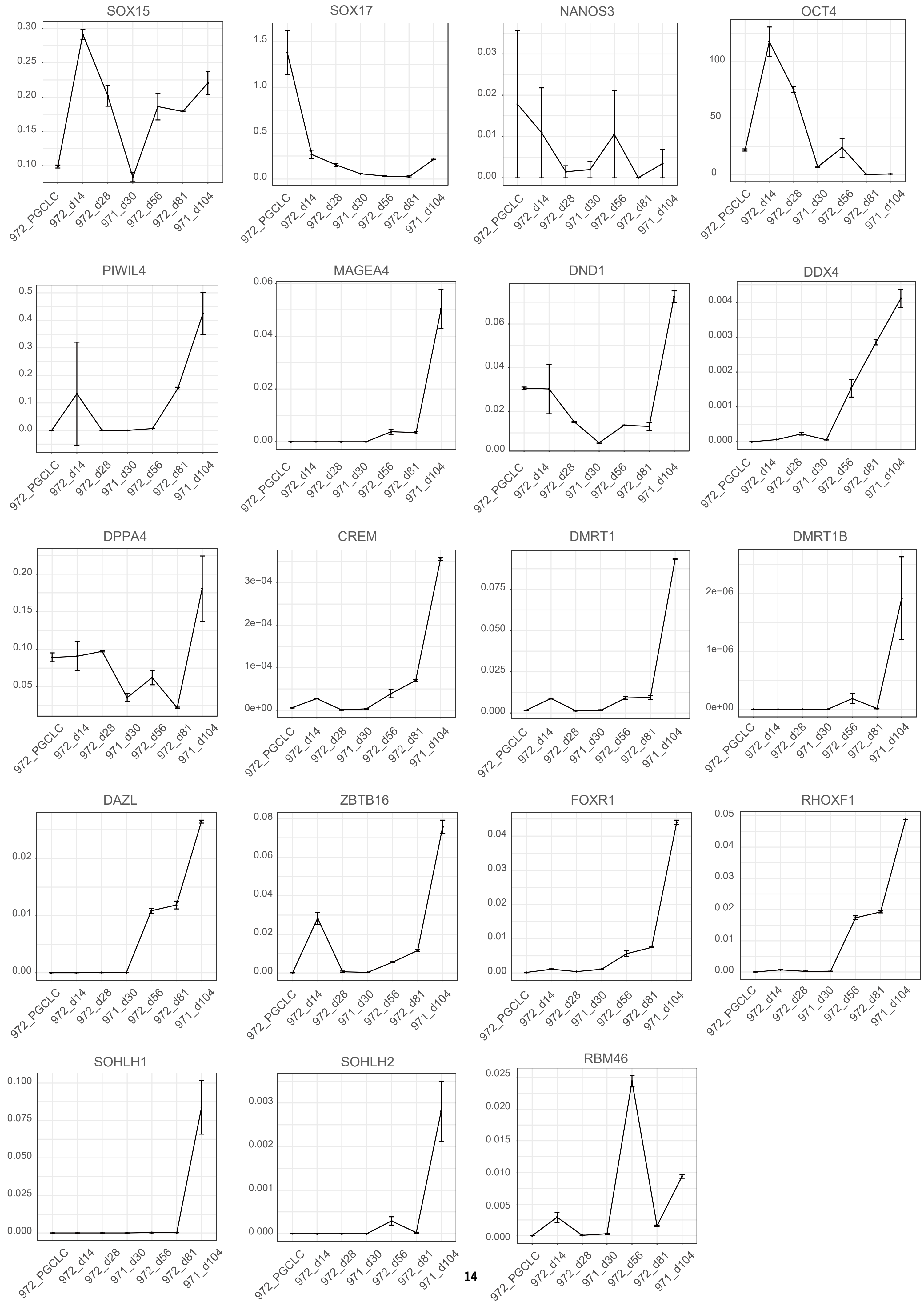
SOX15
SOX17
NANOS3
OCT4
PIWIL4
MAGEA4
DND1
DDX4
DPPA4
CREM
DMRT1
DMRT1B
DAZL
ZBTB16
FOXR1
RHOXF1
SOHLH1
SOHLH2
RBM46
972_PGCLC
972_d14
972_d28
971_d30
972_d56
972_d81
971_d104

Figure S7

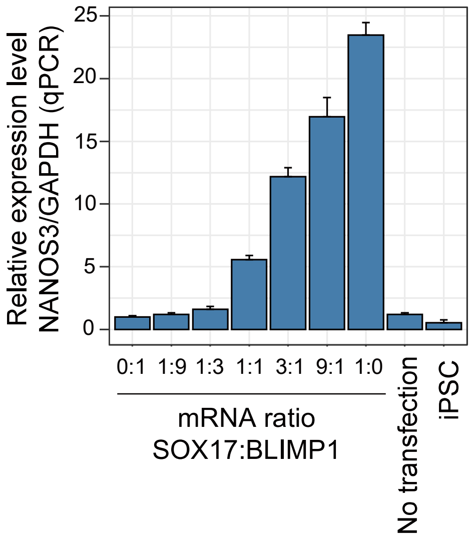
Relative expression level
NANOS3/GAPDH (qPCR)
25
20
15
10
5
0
0:1
1:9
1:3
1:1
3:1
9:1
1:0
No transfection
iPSC
mRNA ratio
SOX17:BLIMP1

Table S1. Sample and cell line information used in this study

| Samples used for 10x library generation | | | | | | | |
|---|---|---|---|---|---|---|---|
| # | Cells or samples (cell line name) | Library type | Animal ID | Sex | Oocyte ID | Sperm ID | Note |
| 1 | iPSC culture (mRNA-STCE) | 10x v3 | I 2965F | F | | | Derived from liver cells. Sox17-T2A-tdTomato knock-in and CAG-EGFP transgene. |
| 2 | PGCLC aggregate (mRNA-STCE) | 10x v3 | I 2965F | F | | | Induced from #1 |
| 3 | E74 ovaries | 10x v2 | | F | I4698F or I5463F | I4213M or YI034M | |
| 4 | E82 ovaries | 10x v3 | | F | I689F | YI810M | |
| 5 | E87 testes | 10x v2 | | M | I4750F or I4962F | I5057M or I764M | |
| 6 | Newborn ovaries | 10x v2 | CM-799 | F | | | |
| 7 | 22 day testes | 10x v2 | I 880M | M | | | |
| 8 | 3yr10mnth testes | 10x v2 | I 6093M | M | | | |

| Samples used for scBS-seq/scRNA-seq library generation | | | | | | | |
|---|---|---|---|---|---|---|---|
| # | Cells or samples | Library type | Animal ID | Sex | Oocyte ID | Sperm ID | Note |
| 9 | iPS/PGCLC (971-STCE) | scBS-seq/ scRNA-seq | I 971M | M | | | Derived from ear cells. |
| 10 | iPS/PGCLC (972-STCE) | scBS-seq/ scRNA-seq | I 972M | M | | | Derived from ear cells. |

Table S2. The list of primer sequences used in this study

| # | Name | |
|---|---|---|
| 1 | Sox17_5UTR | CTCAGAGAGAACCCACCACC ATGAGCAGCCCGGATG |
| 2 | Sox17_R_Asc1 | TTAAC GGCGCGCC TCACACATCAGGATAGTTGCAG |
| 3 | 5UTR_HBA_Not1 | GATCGTACGCGGCCGCTCTTCTGGTCCCCACAGACTCAGAGA GAACCCAC |
| 4 | Sox17 5' arm F | ACGCGTTACGTATCGGATCC TGCAGCACATGCAGGACCAC |
| 5 | Sox17 5' arm R | CTGCCCTCTCCGGATCCCACATCAGGATAGTTGCAGTAATAC AC |
| 6 | Sox17 3' arm F | CGCTAGCGAATTCTAGGATCC GGA GAG CTAAGGAAGTCCTCAG |
| 7 | Sox17 3' arm R | ACGAATTCAGATTCGGATCC AGCCTCTGTAGGCAGGTCAAG |
| 8 | sgRNA-Universal-rev-primer | GTTT GAATTC AAAAAAA GCACCGACTCGGTGCCACTTTTTCAAGTTGATAA CGGACTAGCCTTATTTTAACTTGCTATTTCTAGCTCTAA |
| 9 | pHL_Sox17_1 | CTT GGATCC G AAGCAGTGTTACACACTTCC GTTTTAGA GCTA GAAA TAGCA |
| 10 | pHL_Sox17_2 | CTT GGATCC G ACACACTTCCTGGAGAGCTA GTTTTAGA GCTA GAAA TAGCA |
| 11 | pHL_Sox17_3 | CTT GGATCC G GAGGACTTCCTTAGCTCTCC GTTTTAGA GCTA GAAA TAGCA |
| 12 | Sox17_scr5F | CAAGTCGTGGAAGGCGCTGAC |
| 13 | tdTomatoR1 | GAAGCGCATGAACTCTTTGATGACCTC |
| 14 | Sox17_scr3R | AGGCAAACTCAACGTTGAGGAGTG |
| 15 | human-b-actProR | GACATCTCTTGGGCACTGAG |
| 16 | SOX15-qPCR F | CTACTCGACAGCCTACTTGCC |
| 17 | SOX15-qPCR R | CTGGAGCCTAGGGTCACTCTG |
| 18 | PIWIL4-qPCR F | CAGGTTCCAGTGGAATACCTGTG |
| 19 | PIWIL4-qPCR R | CACATGGTACTGGTATAGCTG |
| 20 | MAGEA4-qPCR F | TGGGAGGAGCTGAGTGTGATG |
| 21 | MAGEA4-qPCR R | CAGAGATACATCTCCAAGTCACTC |
| 22 | DAZL-qPCR F | CTCCGGCTTATTCATCTGTAAACT |
| 23 | DAZL-qPCR R | GATGCACTCTTTTATCCCTGAAGT |
| 24 | DMRT1B-qPCR F | TGCAGTAGGCCCTGAGTACC |
| 25 | DMRT1B-qPCR R | GGTAGTAGCTTGGCTGGAAGTCTC |
| 26 | FOXR1-qPCR F | GCAGAAACTTGCCAGGTATAAACT |
| 27 | FOXR1-qPCR R | GATCCTCCTTCTTCATAAGTCCTG |
| 28 | RHOXF1-qPCR F | GTGTTAACAAGAAGGGAACTTGCT |
| 29 | RHOXF1-qPCR R | ATGTAGAAACTGTCATCTGGATCG |
| 30 | SOHLH1-qPCR F | CTCTGGTGACTGTGGGTCCTA |

| | | |
|---|---|---|
| 31 | SOHLH1-qPCR R | CGTCAACTGTAGAACATTCTCCTT |
| 32 | ZBTB16-qPCR F | ACTATAGGGTGCACACAGGTGAGA |
| 33 | ZBTB16-qPCR R | GAGGTACGTCTTCTCTATCCTCCA |
| 34 | CREM-qPCR F | GTGGAACAATCCAGATTTCTAACC |
| 35 | CREM-qPCR R | GTAGTAGGAGCTCGGATCTGGTAA |
| 36 | DMRT1-qPCR F | ATTCTTACTACCCACCTCCCTCTT |
| 37 | DMRT1-qPCR R | ATGACAGGAGTGACTGTAAAGCTG |
| 38 | SOHLH2-qPCR F | CAGGACATGCAGGTGATATGAC |
| 39 | SOHLH2-qPCR R | CTAGAGATTCAGGGCAGGCAGA |
| 40 | DND1-qPCR F | GGGCAGATCGCTCTGCTC |
| 41 | DND1-qPCR R | CTCTCCACAGAGGTGTGATTG |
| 42 | SOX17-qPCR F | CCGAGCTGAGCAAGATGCTG |
| 43 | SOX17-qPCR R | GTGGTCCTGCATGTGCTGCAC |
| 44 | NANOS3-qPCR F | CTTCTGCCCACTCACTGGACAG |
| 45 | NANOS3-qPCR R | CTCAGACTTCCCGGCACCTCTG |
| 46 | DDX4-qPCR F | AAGTATTAACAGATGCTCAACAGGATGT |
| 47 | DDX4-qPCR R | TGAAGCCAGGAATGTATGCACTA |
| 48 | DPPA4-qPCR F | TGGGTAAGCAAAGGCACACAG |
| 49 | DPPA4-qPCR R | CTGGTGTCAGCAACTAAAGCTAAGCAC |
| 50 | GAPDH-qPCR F | TGCTGGCGCTGAGTATGTG |
| 51 | GAPDH-qPCR R | AGCCCCAGCCTTCTCCAT |
| 52 | CB16UMI12-RT primer1 | TCA GAC GTG TGC TCT TCC GAT CTA ATC GGT GTT GAT TCG NNN NNN NNN NNN TTT TTT TTT TTT TTT TTT TTT TTT T |
| 53 | BC16UMI12-RT primer2 | TCA GAC GTG TGC TCT TCC GAT CTA CAG CTA CAC GTG AGA NNN NNN NNN NNN TTT TTT TTT TTT TTT TTT TTT TTT T |
| 54 | BC16UMI12-RT primer3 | TCA GAC GTG TGC TCT TCC GAT CTA GCG TAT AGA CGA CGT NNN NNN NNN NNN TTT TTT TTT TTT TTT TTT TTT TTT T |
| 55 | BC16UMI12-RT primer4 | TCA GAC GTG TGC TCT TCC GAT CTA TCA TCT GTA GCG TAG NNN NNN NNN NNN TTT TTT TTT TTT TTT TTT TTT TTT T |
| 56 | BC16UMI12-RT primer5 | TCA GAC GTG TGC TCT TCC GAT CTC ACA TAG TCG CAC TCT NNN NNN NNN NNN TTT TTT TTT TTT TTT TTT TTT TTT T |
| 57 | BC16UMI12-RT primer6 | TCA GAC GTG TGC TCT TCC GAT CTC CTA CAC TCT ACT ATC NNN NNN NNN NNN TTT TTT TTT TTT TTT TTT TTT TTT T |
| 58 | BC16UMI12-RT primer7 | TCA GAC GTG TGC TCT TCC GAT CTC GAG CAC AGA TAG CAT NNN NNN NNN NNN TTT TTT TTT TTT TTT TTT TTT TTT T |
| 59 | BC16UMI12-RT primer8 | TCA GAC GTG TGC TCT TCC GAT CTC TGA AGT AGT ATT GGA NNN NNN NNN NNN TTT TTT TTT TTT TTT TTT TTT TTT T |

| | | |
|---|---|---|
| 60 | BC16UMI12-RT primer9 | TCA GAC GTG TGC TCT TCC GAT CTG ACA CGC TCA GTC AGT NNN NNN NNN NNN TTT TTT TTT TTT TTT TTT TTT TTT T |
| 61 | BC16UMI12-RT primer10 | TCA GAC GTG TGC TCT TCC GAT CTG CAA TCA CAA TGT TGC NNN NNN NNN NNN TTT TTT TTT TTT TTT TTT TTT TTT T |
| 62 | BC16UMI12-RT primer11 | TCA GAC GTG TGC TCT TCC GAT CTG GTG CGT AGG TGC ACA NNN NNN NNN NNN TTT TTT TTT TTT TTT TTT TTT TTT T |
| 63 | BC16UMI12-RT primer12 | TCA GAC GTG TGC TCT TCC GAT CTG TTC TCG TCT GCT GTC NNN NNN NNN NNN TTT TTT TTT TTT TTT TTT TTT TTT T |
| 64 | BC16UMI12-RT primer13 | TCA GAC GTG TGC TCT TCC GAT CTT ACT CAT CAC AGT CGC NNN NNN NNN NNN TTT TTT TTT TTT TTT TTT TTT TTT T |
| 65 | BC16UMI12-RT primer14 | TCA GAC GTG TGC TCT TCC GAT CTT CGG TAA TCA CGC ATA NNN NNN NNN NNN TTT TTT TTT TTT TTT TTT TTT TTT T |
| 66 | BC16UMI12-RT primer15 | TCA GAC GTG TGC TCT TCC GAT CTT GTC CCA GTT TGG CGC NNN NNN NNN NNN TTT TTT TTT TTT TTT TTT TTT TTT T |
| 67 | BC16UMI12-RT primer16 | TCA GAC GTG TGC TCT TCC GAT CTT TGA CTT GTA CTC GCG NNN NNN NNN NNN TTT TTT TTT TTT TTT TTT TTT TTT T |
| 68 | TSO primer | /5Me-isodC//iisodG//iMe-isodC/AAGCAGTGGTATCAACGCAGAGTACATrGrG+G |
| 69 | 3' Anchoerd primer | GTGACTGGAGTTCAGACGTGTGCTCTTCCGATC |
| 70 | ISPCR primer | AAGCAGTGGTATCAACGCAGAGT |
| 71 | scBS-P5-N9-oligo1 | CTACACGACGCTCTTCCGATCTNNNNNNNNN |
| 72 | scBS-oligo2 | GACTGGAGTTCAGACGTGTGCTCTTCCGATCTNNNNNNNNN |
| 73 | Biotin-index-primer103 | Biotin/CAAGCAGAAGACGGCATACGAGATAGAGTAGTGACTGGAGTTCAGACGTGTGCTCTTCCGATC |
| 74 | Biotin-index-primer104 | Biotin/CAAGCAGAAGACGGCATACGAGATAGTCCAGTGACTGGAGTTCAGACGTGTGCTCTTCCGATC |
| 75 | Biotin-index-primer105 | Biotin/CAAGCAGAAGACGGCATACGAGATGCCAATGTGACTGGAGTTCAGACGTGTGCTCTTCCGATC |
| 76 | Biotin-index-primer106 | Biotin/CAAGCAGAAGACGGCATACGAGATCTTGTAGTGACTGGAGTTCAGACGTGTGCTCTTCCGATC |
| 77 | QP2 | CAAGCAGAAGACGGCATACGA |
| 78 | Modified_P5 primer | AATGATACGGCGACCACCGAGATCTACAC[index]ACACTCTTTCCCTACACGAC |
